# Diversity and function of fungi associated with the fungivorous millipede, *Brachycybe lecontii*

**DOI:** 10.1101/515304

**Authors:** Angie M. Macias, Paul E. Marek, Ember M. Morrissey, Michael S. Brewer, Dylan P.G. Short, Cameron M. Stauder, Kristen L. Wickert, Matthew C. Berger, Amy M. Metheny, Jason E. Stajich, Greg Boyce, Rita V. M. Rio, Daniel G. Panaccione, Victoria Wong, Tappey H. Jones, Matt T. Kasson

## Abstract

Fungivorous millipedes (subterclass Colobognatha) likely represent some of the earliest known mycophagous terrestrial arthropods, yet their fungal partners remain elusive. Here we describe relationships between fungi and the fungivorous millipede, *Brachycybe lecontii*. Their fungal community is surprisingly diverse with 176 genera, 39 orders, and four phyla and includes several undescribed species. Of particular interest are twelve genera conserved across wood substrates and millipede clades that comprise the core fungal community of *B. lecontii.* Wood decay fungi, long speculated to serve as the primary food source for *Brachycybe* species, were absent from this core assemblage and proved lethal to millipedes in pathogenicity assays while entomopathogenic Hypocreales were more common in the core but had little effect on millipede health. This study represents the first survey of fungal communities associated with any colobognath millipede, and these results offer a glimpse into the complexity of millipede fungal communities.

## 1. Introduction

The Class Diplopoda, known colloquially as millipedes, represent some of the earliest known terrestrial animals, dating back to the early Devonian period *ca.* 412 million years ago (Suarez *et al.* 2017, Wilson & Anderson 2004). These early representatives were detritivores and likely played a role in early soil formation and the development of terrestrial nutrient cycling (Bonkowski *et al.* 1998, Lawrence & Samways 2003). Detritivorous millipedes continue to play a pivotal role in ecosystem processes, though herbivorous (Marek *et al.* 2012), carnivorous (Srivastava & Srivastava 1967), and fungivorous diets also exist among extant millipedes (Brewer *et al.* 2012, Marek *et al.* 2012).

Most fungivorous millipedes belong to the subterclass Colobognatha, which diverged from detritus-feeding millipedes 200-300 million years ago and possess a primitive trunk-ring architecture composed of a free sternum and/or pleurites (Brewer & Bond 2013, Rodriguez *et al.* 2018). However, these millipedes are characterized by derived rudimentary mouthparts adapted for feeding exclusively on succulent tissues such as fungi (Hopkin & Read 1992; Wilson & Anderson 2004, Wong 2018). Some taxa possess beaks composed of elongate mouthparts encompassing a fused labrum, gnathochilarium, and stylet like mandibles (Read & Enghoff 2018). Members of the Colobognatha are among the most understudied groups in Diplopoda despite their wide geographic distribution and ubiquity in natural history collections (Hoffman 1980, Manton 1961, Read & Enghoff 2009, Shorter *et al.* 2018). The ancient association between millipedes and fungi raises fascinating questions about interactions in early terrestrial ecosystems, and the possible role of fungi in diplopod success.

Most published records of fungal-millipede interactions are cases where millipedes graze on fungi in the environment (Lilleskov & Bruns 2005, Bultman & Mathews 1996) or where parasitic fungi infect millipedes (Hodge *et al.* 2017, Kudo *et al.* 2011). Among the most studied millipede fungal associates are specialist ectoparasitic fungi in the Laboulbeniales (Santamaria *et al.* 2014, Enghoff & Santamaria 2015, Reboleira *et al.* 2018) and obligate arthropod gut-associated trichomycetes (Wright 1979). However, in none of these interactions does the millipede strictly depend on fungi for survival as it seemingly does in the fungivorous millipede *B. lecontii* (Diplopoda: Platydesmida: Andrognathidae).

*Brachycybe lecontii* are most frequently found in multigenerational aggregations in decaying wood with visible fungal growth (Gardner 1975, Shelley *et al.* 2005). The known geographic range of *B. lecontii* extends across 13 U.S. states from eastern Oklahoma to western South Carolina, south to Louisiana, and north to southern West Virginia (Shelley *et al.* 2005, Brewer *et al.* 2012). Within its known range, *B. lecontii* is divided into at least 4 clades that are geographically separated and may represent independent cryptic species (Brewer *et al.* 2012).

Historically, only one study reported *Brachycybe* feeding on an identified fungus, an unknown species of *Peniophora* (Russulales) (Gardner 1975). However, observations of *Brachycybe* species interacting with various fungi (n = 65) from community science websites such as Bugguide.net and iNaturalist.org shows that the fungal communities associated with this genus are more diverse than have been formally described (Supplemental Table 1). Recently, several basidiomycete Polyporales have been confirmed directly from *B. lecontii* and from *B.* lecontii-associated wood (Kasson *et al.* 2016). Given the discovery that Polyporales have helped facilitate the evolution of large, communal colonies with overlapping generations in other arthropods (You *et al.* 2015, Simmons *et al.* 2016, Kasson *et al.* 2016), many interesting questions are raised regarding *Brachycybe* colonies and their association with Polyporales and allied fungi.

In an attempt to uncover which fungi, if any, are consistently associated with *B. lecontii,* this study surveys fungal associates of *B. lecontii* across its known geographic range using culture-based approaches. The use of nuclear ribosomal internal transcribed spacer (ITS) barcoding on collected isolates allowed for fine-scale identification and examination of culturable fungal communities. With a primary understanding of these fungal communities, we assessed diversity and applied a network analysis to determine the relationship between genetically and geographically distinct *B. lecontii* populations, their wood substrates, and associated fungal genera.

## 2. Materials and methods

### 2.1. Collection sites & field methods

Millipede collection sites were primarily identified through Brewer *et al.* (2012) and Gardner (1975), and additional sites were identified based on the known range of *B. lecontii.* Sampling was targeted to collect millipedes from all four *B. lecontii* clades. Based on previous work by Brewer (2012), individual sites were expected to contain millipedes from a single clade, with no syntopy. In total, 20 sites were sampled, with 18 yielding colonies and solitary individuals, and 2 yielding individuals only. These sites were located in Arkansas, Georgia, Oklahoma, South Carolina, Tennessee, Virginia, and West Virginia (Table 1).

**Table 1.** *Brachycybe lecontii* collection sites.

| Site | Collection reference | Millipede<br>clade | Level 4<br>ecoregion | # millipedes<br>sampled |
| --- | --- | --- | --- | --- |
| AR1 | Gardner 1975 | 4 | 37A | 6 |
| AR2 | ----- | 4 | 36B | 6 |
| AR3 | ----- | 4 | 36C | 13 |
| AR4 | Brewer *et al.* 2012 | 4 | 37A | 22 |
| AR5 | ----- | 4 | 36B | 14 |
| AR6 | ----- | 4 | 36B | 12 |
| GA1 | Gardner 1975 | 1 | 66D | 9 |
| OK1 | Brewer *et al.* 2012 | 4 | 36D | 14 |
| SC1 | Brewer *et al.* 2012,<br>Gardner 1975 | 1 | 66D | 22 |
| SC2 | Gardner 1975 | 1 | 45E | 4 |
| TN1 | Gardner 1975 | 3 | 69D | 42 |
| VA1 | Brewer *et al.* 2012,<br>Gardner 1975 | 3 | 69D | 4 |
| VA2 | ----- | 3 | 67I | 8 |
| VA3 | Gardner 1975 | 3 | 69D | 15 |
| VA4 | ----- | 1 | 67H | 5 |
| WV1 | Brewer *et al.* 2012,<br>Gardner 1975 | 3 | 69D | 29 |
| WV2 | ----- | 3 | 69D | 24 |
| WV3 | ----- | 3 | 69D | 4 |
| WV4 | ----- | 3 | 69D | 12 |
| WV5 | ----- | 3 | 69D | 36 |

At each site, decaying logs on the forest floor were overturned, examined, and replaced until colonies of *B. lecontii* were located. Colonies are defined as groups of two or more individuals, and were typically found on or near resupinate fungi covering the underside of the logs. When suitable colonies were found, individuals from single colonies were placed together in 25-ml sterile collection vials, often with a piece of the fungus-colonized wood they were on, and stored in a cooler until processing. In addition, cross-sections of logs from which colonies were collected were sampled for wood substrate identification.

### 2.2. Millipede processing & isolate collection

All millipedes were maintained at 4°C until processing, which typically occurred within three days of initial collection. After surface sterilization in 70% ethanol, individuals were sexed, sectioned with a sterilized scalpel to remove tail and gonopod sections (Macias 2017). Tail portions were preserved in 95% ethanol for millipede genotyping using custom markers previously described by Brewer et al. (2012). Gonopods were also preserved from males to permit anatomical study. The remainder of the millipede was macerated in 500 μl of sterile distilled water, and a 50-μl sample was spread on glucose yeast extract agar (GYEA) amended with streptomycin sulfate and tetracycline hydrochloride antibiotics to isolate fungi (Macias 2017). Cultures were sealed with parafilm and incubated at ambient conditions until growth was observed. Each colony-forming unit (CFU) was categorized by morphotype, counted, and recorded. One representative of each morphotype from each plate was retained and assigned an isolate number. Culture plates were retained for up to three weeks to ensure that slow-growing fungi were counted and sampled. Depending on how rapidly fungi grew in pure culture, isolates were either grown on potato dextrose broth (PDB; BD and Co., Franklin Lakes, NJ, USA) prior to DNA extraction, or mycelium was scraped directly from plates. DNA was extracted from all isolates using a Wizard kit (Promega, Madison, WI, USA). DNA was suspended in 75 ml of Tris-EDTA (TE) buffer preheated to 65°C. For long-term storage, isolates were kept on potato dextrose agar slants (PDA; BD and Co., Franklin Lakes, NJ, USA) at 4°C.

Wood samples were dried at room temperature for several weeks and sanded using an orbital sander equipped with 220-grit paper. Identifications were made by examining wood anatomy in cross section, based on descriptions by Panshin and de Zeeuw (1980).

### 2.3 Isolate identification

Isolates were identified using the universal fungal barcoding gene, the ribosomal internal transcribed spacer region (ITS), which includes ITS1, 5.8S, and ITS2 (Schoch *et al.* 2012). Primers used in this study were obtained from Integrated DNA Technologies (IDT, Coralville, IA, USA). PCR was conducted using primers ITS4 and ITS5 (White *et al.* 1990), following the protocol in Macias (2017).

PCR products were visualized via gel electrophoresis on a 1.5% w/v agarose (Amresco, Solon, OH, USA) gel with 0.5% Tris-Borate-EDTA buffer (Amresco, Solon, OH, USA). SYBR Gold (Invitrogen, Grand Island, NY, USA) was used as the nucleic acid stain, and bands were visualized on a UV transilluminator (Syngene, Frederick, MD, USA). PCR products were purified using ExoSAP-IT (Affymetrix, Santa Clara, CA). Products were Sanger sequenced with the same primers used for PCR (Eurofins, Huntsville, AL, USA). Resulting sequences were clipped using the “Clip ends” function in CodonCode Aligner v 5.1.5 and searched in the NCBI GenBank BLASTn database (Altschul *et al.* 1990) and best-match identifications recorded for each isolate.

### 2.4. Identification of new species

Fungal isolates were considered to represent a putative new species if three or more identical sequences were recovered with identical low percentage (threshold ≤ 95%) BLASTn matches. The large subunit of the ribosomal ITS region (LSU) was also sequenced using primers LR0R and LR5 (Vilgalys and Hester 1990) for each putative new species. PCR conditions were as described in Macias (2017). PCR products were visualized, purified, and sequenced as above.

In the following analyses, the default parameters of each software package were used unless otherwise noted. Two putative new species were confirmed phylogenetically by constructing ITS+LSU concatenated phylogenetic trees for each new species and its known relatives based on a combination of BLAST matches and previously published literature. MEGA7 v. 7.0.16 (Kumar *et al.* 2016) was used to align sequences (CLUSTAL-W, Larkin *et al.* 2007), select a best-fit model for estimating phylogeny, and construct maximum likelihood (ML) and maximum parsimony (MP) trees for each putative new species. For ML and MP analyses, 1000 bootstrap replicates were used. The alignments used default parameters. Initial alignments were trimmed such that all positions with less than 95% site coverage were eliminated. For maximum likelihood analyses, the Tamura 3-parameter substitution model with a gamma distribution (T92+G) was used (Tamura 1992). Both maximum parsimony analyses used the subtree-pruning-regrafting algorithm (Nei and Kumar 2000). Bayesian (BI) trees were constructed using Mr. Bayes v. 3.2.5 (Ronquist *et al.* 2012). Three hot and one cold independent MCMC chains were run simultaneously for 1,000,000 generations, and the first 25% were discarded as a burn-in. The average standard deviation of split frequencies statistic was checked to ensure convergence between chains (Ronquist *et al.* 2012) and was <0.01. Final parameter values and a final consensus tree were generated used the MrBayes “sump” and “sumt” commands respectively. The ML tree was preferred and support for its relationships in the other analyses was determined. Reference sequences used in each phylogenetic analysis are listed in Supplemental Tables 2 and 3, respectively.

### 2.5. Community and diversity analyses

Community and diversity analyses were used to answer two questions. 1) Are *B. lecontii* fungal communities conserved across millipede clade, millipede sex, wood substrate, and/or ecoregion? 2) What genera comprise the core fungal community associated with *B. lecontii*?

The effects of *B. lecontii* clade, sex, wood substrate, and ecoregion on the fungal community composition were analyzed by perMANOVA using the vegan package (Oksanen *et al.* 2018) in R version 3.4.3 (R Core Team 2017). Multilevel pairwise comparisons were performed using the pairwiseAdonis package (Arbizu 2017). Isolates where the fungal order or wood substrate were not identified were removed from the analysis. Additionally, isolates recovered from millipedes not part of a colony were removed. Ecoregion 45 was deleted from the analysis because its variance was significantly different from the other ecoregions (checked using function betadisper in vegan). Pairwise comparisons were only made for groups with 20 or more millipedes sampled, and in cases where more than one pairwise comparison was made, Bonferroni-corrected p-values are reported.

In addition, diversity indices were used to provide information about rarity and commonness of genera in the fungal community of *B. lecontii,* by site. Three alpha diversity indices were chosen according to recommendations laid out in Morris *et al.* (2014): number of genera present, Shannon’s diversity index, and Shannon’s equitability (evenness) index (formulae from Begon *et al.* 1990). The relationship between sample size and the three diversity metrics was examined using Spearman rank correlations.

A co-occurrence network was constructed using Gephi (Bastian *et al.* 2009) for fungal isolates obtained from *B. lecontii* at the genus level. Betweenness-centrality was used to measure relative contribution of each node (single fungal genus) to connectivity across the whole network. High betweenness-centrality values are typically associated with nodes located in the core of the network (Greenblum *et al.* 2011), which in this system are defined as fungal genera with multiple edges connecting multiple *B. lecontii* clades and multiple wood substrates. Low betweenness-centrality values indicate fungal genera with a more peripheral location in the network, with fewer edges connecting clades and wood substrates (Greenblum *et al.* 2011).

### 2.6. Pathogenicity testing

Twenty-one isolates (Table 2) representing the diversity of all collected isolates were initially chosen for live plating pathogenicity assay (hereafter referred to as “full-diversity assay”) with *Brachycybe lecontii.* A second assay (hereafter referred to as “Polyporales assay”) using nineteen isolates in the Polyporales (Table 3) were tested in a separate experiment. Three isolates from the full-diversity assay were re-tested in the second assay, for a total of 37 isolates used between both assays.

**Table 2.** Results of pathogenicity assays using a representative set of fungal community members isolated from *B. lecontii.* HPP = Hours post-plating. Asterisks denote significantly faster and greater mortality in a treatment, compared to the negative control: * = p<0.05, ** = p<0.01, *** = p<0.001. Both Log-rank P and Wilcoxon P are Bonferroni-corrected. † Denotes isolates tested again in the Polyporales pathogenicity assay.

| Name | Order | Isolate | Time to 50% mortality | % dead at 7 days HPP | Log-rank P | Wilcoxon P |
| --- | --- | --- | --- | --- | --- | --- |
| Negative control | ----- | ----- | ----- | 0% | ----- | ----- |
| *Pestalotiopsis microspora* | Amphisphaeriales | BC0630 | ----- | 20% | ----- | ----- |
| *Chaetosphaeria myriocarpa* | Chaetosphaeriales | BC0320 | ----- | 0% | ----- | ----- |
| *Capronia dactylotricha* | Chaetothyriales | BC1244 | ----- | 0% | ----- | ----- |
| *Fonsecaea* sp. | Chaetothyriales | BC1147 | ----- | 0% | ----- | ----- |
| *Phialophora americana* | Chaetothyriales | BC1193 | ----- | 7% | ----- | ----- |
| *Lecanicillium attenuatum* | Hypocreales | BC0678 | ----- | 0% | ----- | ----- |
| *Metarhizium flavoviride* | Hypocreales | BC1163 | 136 HPP | 87% | <.0005*** | <.0005*** |
| *Pochonia bulbillosa* | Hypocreales | BC0029 | ----- | 0% | ----- | ----- |
| *Trichoderma viride* | Hypocreales | BC1216 | ----- | 7% | ----- | ----- |
| *Verticillium insectorum* | Hypocreales | BC0482 | ----- | 0% | ----- | ----- |
| *Mortierella* aff. *ambigua* | Mortierellales | BC1150 | ----- | 0% | ----- | ----- |
| *Mortierella* sp. | Mortierellales | BC0530 | ----- | 27% | 0.173 | 0.175 |
| aff. *Apophysomyces* sp. | Mucorales | BC1015 | ----- | 0% | ----- | ----- |
| *Mucor abundans* | Mucorales | BC1010 | ----- | 0% | ----- | ----- |
| *Ramichloridium anceps* | Mycosphaerellales | BC0329 | ----- | 0% | ----- | ----- |
| *Bjerkandera adusta*† | Polyporales | BC0310 | 36 HPP | 80% | <.0005*** | <.0005*** |
| *Irpex lacteus*† | Polyporales | BC0523 | 72 HPP | 47% | 0.014* | 0.0155* |
| *Trametopsis cervina*† | Polyporales | BC1143 | 36 HPP | 100% | <.0005*** | <.0005*** |
| *Umbelopsis angularis* | Umbelopsidales | BC0529 | ----- | 13% | ----- | ----- |
| *Umbelopsis isabellina* | Umbelopsidales | BC1290 | ----- | 0% | ----- | ----- |
| *Umbelopsis ramanniana* | Umbelopsidales | BC1028 | ----- | 0% | ----- | ----- |
| Overall | ----- | ----- | ----- | ----- | <.0001*** | <.0001*** |

**Table 3.** Results of pathogenicity assays using a representative set of Polyporales isolated from B. lecontii. HPP = Hours post-plating. Asterisks denote significantly faster and greater mortality in a treatment, compared to the negative control: * = p<0.05, ** = p<0.01, *** = p<0.001. Both Log-rank P and Wilcoxon P are Bonferroni-corrected. † Denotes isolates used in initial pathogenicity assay.

| Name | Order | Isolate | Time to<br>50%<br>mortality | % dead at 7<br>days HPP | Log-rank<br>P | Wilcoxon<br>P |
| --- | --- | --- | --- | --- | --- | --- |
| Negative control | ----- | ----- | ----- | 20% | ----- | ----- |
| *Bjerkandera adusta*† | Polyporales | BC0310 | 36 HPP | 50% | 0.8484 | 0.4907 |
| *Ceriporia lacerata* | Polyporales | BC1158 | ----- | 10% | ----- | ----- |
| *Ceriporiopsis gilvescens* | Polyporales | BC0174 | ----- | 0% | ----- | ----- |
| *Gloeoporus pannocinctus* | Polyporales | BC0042 | 12 HPP | 100% | <.0007*** | <.0007*** |
| *Irpex lacteus* | Polyporales | BC1038 | ----- | 30% | 1.0000 | 1.0000 |
| *Irpex lacteus*† | Polyporales | BC0523 | 72 HPP | 50% | 0.7119 | 0.5033 |
| *Junghuhnia nitida* | Polyporales | BC0528 | ----- | 10% | ----- | ----- |
| *Phanerochaete*<br>*cumulodentata* | Polyporales | BC0709 | ----- | 0% | ----- | ----- |
| *Phanerochaete sordida* | Polyporales | BC0691 | 84 HPP | 60% | 0.2352 | 0.1603 |
| *Phlebia acerina* | Polyporales | BC1331 | ----- | 0% | ----- | ----- |
| *Phlebia fuscoatra* | Polyporales | BC1014 | ----- | 0% | ----- | ----- |
| *Phlebia livida* | Polyporales | BC0629 | ----- | 0% | ----- | ----- |
| *Phlebia subserialis* | Polyporales | BC0054 | ----- | 0% | ----- | ----- |
| *Phlebiopsis flavidoalba* | Polyporales | BC1426 | ----- | 10% | ----- | ----- |
| *Phlebiopsis gigantea* | Polyporales | BC1049 | ----- | 0% | ----- | ----- |
| *Scopuloides rimosa* | Polyporales | BC1046 | ----- | 0% | ----- | ----- |
| *Trametopsis cervina*† | Polyporales | BC1143 | 36 HPP | 100% | <.0007*** | <.0007*** |
| *Trametopsis cervina* | Polyporales | BC0494 | 36 HPP | 100% | <.0007*** | <.0007*** |
| *Peniophora pithya* | Russulales | BC1467 | ----- | 0% | ----- | ----- |
| Overall | ----- | ----- | ----- | ----- | <.0001*** | <.0001*** |

Isolates were grown on GYEA and scraped to generate inoculum suspensions in sterile water. An aliquot of suspension was spread onto fresh GYEA plates. After all plates were covered by fungal growth (~3 weeks), the millipedes were introduced for 7-day pathogenicity trials. Five individuals were placed on each plate. In the full-diversity assay, 15 millipedes were used for each treatment, while 10 were used in the Polyporales assay. For a negative control, millipedes were placed on sterile GYEA plates that were changed each time contaminating fungal growth was observed. These plates required replacement due to inadvertent inoculation by the millipede’s phoretic contaminants and gut microbes. Observations were made every 12 hours for the first 36 hours and then every 4 hours for an additional 108 hours until 7 days were complete. Mortality was assessed by failure of millipedes to move in response to external stimuli (Panaccione & Arnold 2017). At the end of the assay, samples of deceased individuals were preserved for chemical analyses. Surviving millipedes were returned to laboratory colonies for future studies.

Statistical analysis of survivorship was performed using the “Survival / Reliability” function in JMP 13.1.0 (SAS Institute Inc., Cary, NC). Post-hoc pairwise comparisons to the control treatment were performed for treatments that reached at least 25% mortality by the end of the trials (five treatments in the full-diversity assay, and seven in the Polyporales assay). Both log-rank and Wilcoxon tests were used. Log-rank tests score mortality at all time points evenly, while Wilcoxon tests score early mortality more heavily. For pairwise comparisons, Bonferroni corrections were applied such that the P-value reported by the analysis was multiplied by the number of comparisons made in each experiment.

## 3. Results and discussion

### 3.1. Diversity and community structure

A total of 301 millipedes were collected from 3 of 4 known *B. lecontii* clades (Brewer *et al.* 2012) and from 20 sites across 7 states (Table 1). Our study recovered 102 males and 146 mature females. Most millipedes were engaged in feeding behavior, with their heads buried in fungus growing on the log (Figure 1).

**Figure 1.**
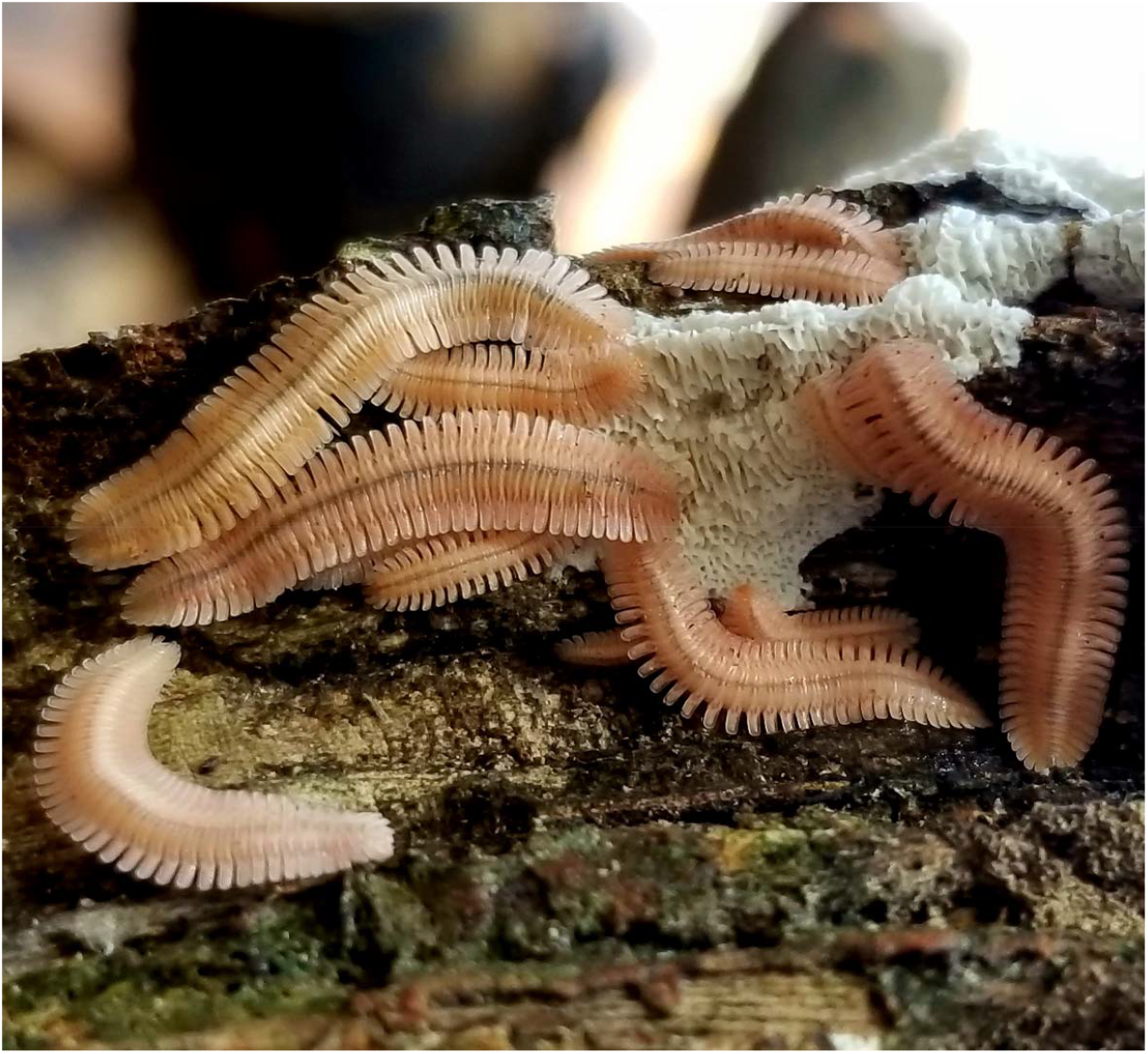
*Brachycybe lecontii* colony feeding on the white-rot fungus *Irpex lacteus* (Basidiomycota: Polyporales).

*Brachycybe lecontii* was found to associate with a large and diverse community of fungi, including at least 176 genera in 39 fungal orders from four phyla (Supplemental Table 4). A majority of these fungi (59%) were members of Ascomycota. Of all the genera of fungi found in this study, 40% were represented by a single isolate, and only 13% had 10 or more isolates. The most common order was the Hypocreales, containing 26% of all isolates resolved to at least order. The five next most common orders were the Polyporales (9% of all isolates), Chaetothyriales (8%), Xylariales (6%), Capnodiales (5%), and Eurotiales (5%). All other orders contained fewer than 50 isolates (<5%).

**Table 4.**
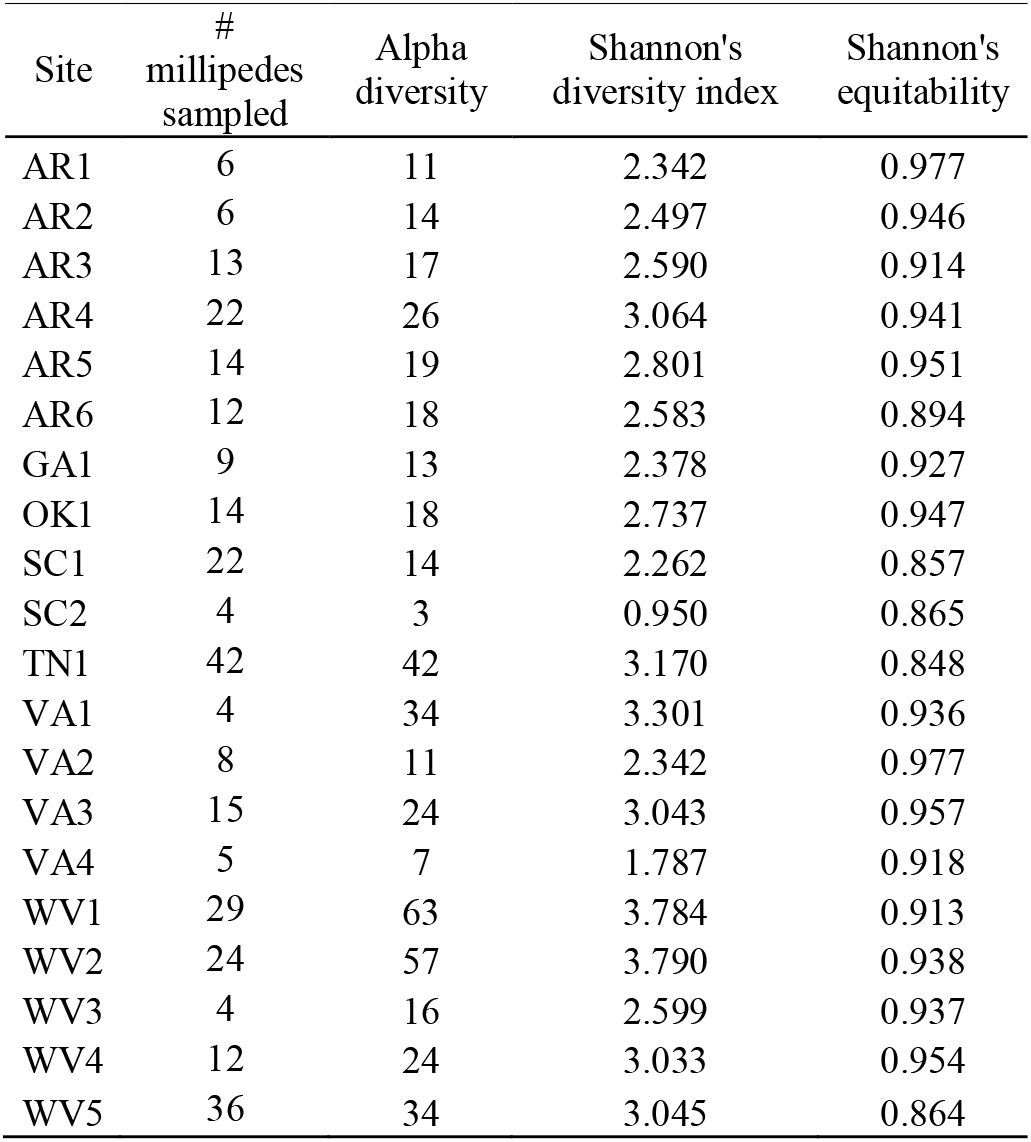
Collection information and genus-level diversity indices for each site.

Alpha diversity was assessed at the genus level by millipede clade, wood substrate, and site. Clade 1 included 31 fungal genera, clade 3 included 156, and clade 4 included 69. The number of fungal genera obtained from each wood substrate were as follows: *Liriodendron* (114 genera), *Quercus* (74), *Betula* (54), *Carya* (45), *Fagus* (33), *Ulmus* (19), *Acer* (18), *Pinus* (11), *Carpinus* (10), and *Fraxinus* (10). However, the number of millipedes sampled for Clade 1, and all wood substrates except *Quercus* and *Liriodendron*, are likely not sufficient to capture the full diversity of the communities with culture-based methods (n<20).

At each site, fungal alpha diversity varied from 3 genera at SC2 to 63 genera at WV1 with a mean of 23 per site (Table 4). Shannon’s diversity index and Shannon’s equitability were also calculated for each site. Shannon’s diversity index ranged from 0.95 in SC2 to 3.79 in WV2. Shannon’s equitability ranged from 0.977 in AR1 and VA2 to 0.848 in TN1 (Table 4). Sites with the five highest Shannon’s diversity index values did not overlap with sites with the five highest site equitability values (Table 4). Three diversity metrics were found to be correlated with sample size (alpha diversity r = 0.70, Shannon’s diversity index r = 0.58, Shannon’s equitability index r = -0.28), indicating that many more millipedes would be needed to capture the full fungal diversity using culture-based methods. The wide ranges of the three diversity metrics across sites raises questions about functional redundancies in the fungal communities.

To statistically explore relationships between fungal community composition and millipede sex, millipede clade, wood substrate, and ecoregion, perMANOVAs and pairwise multilevel comparisons were used. No relationship was found between millipede sex and fungal community composition (p = 0.353, R^2^ = 0.005), but relationships were found for millipede clade (p = 0.045, R^2^ = 0.006), wood substrate (p = 0.002, R^2^ = 0.049), and ecoregion (p = 0.001, R^2^ = 0.012). However, effect size is very small for each of these factors, indicating that while there are significant relationships between these factors and the fungal community, the strength of those relationships is weak. Pairwise multilevel corrections indicate that there are significant differences between the fungal communities of Clade 1 and 3 (p = 0.009, R^2^ = 0.015) and 4 and 3 (p = 0.003, R^2^ = 0.015), but not 1 and 4 (p = 0.096, R^2^ = 0.014). For wood substrate, only the fungal communities of *Liriodendron* and *Quercus* were compared (see Methods), and they were significantly different (p = 0.045, R^2^ = 0.015). For ecoregion, there are significant differences between the fungal communities of ecoregions 36 and 39 (p = 0.018, R^2^ = 0.014) and 66 and 69 (p = 0.006, R^2^ = 0.018), but not 36 and 37 (p = 1.000, R^2^ = 0.007), 36 and 66 (p = 0.084, R^2^ = 0.025), 37 and 66 (p = 0.462, R^2^ = 0.029), or 37 and 69 (p = 0.300, R^2^ = 0.010). For all of these pairwise comparisons, the effect size was small. Increased sampling depth should help clarify whether any of these factors truly impact the fungal community composition.

A network analysis and betweenness-centrality scores were used to examine how the structure of the fungal community is affected by different millipede clades and wood substrates (Figure 2). As a whole, community structure was heterogeneous across millipede clades and wood substrates. However, some genera of fungi were consistently associated with most clades and wood substrates, as indicated by their betweenness-centrality scores. Twelve fungal genera showed high connectivity across the whole network (betweenness-centrality values > 0.5) (Figure 2). These included 1) *Phialophora* (1.55), 2) *Ramichloridium* (1.44), 3) *Mortierella* (1.28), 4) *Trichoderma* (1.03), 5) *Mucor* (1.02), 6) *Verticillium* (0.90), 7) *Phanerochaete* (0.89), 8) *Fonsecaea* (0.84), 9) *Penicillium* (0.75), 10) *Umbelopsis* (0.73), 11) *Cosmospora* (0.68), and 12) *Xylaria* (0.63). All other fungal genera fell below a 0.5 threshold, including 144 genera with betweenness-centrality values of 0.0, indicating low presence across millipede communities.

**Figure 2.**
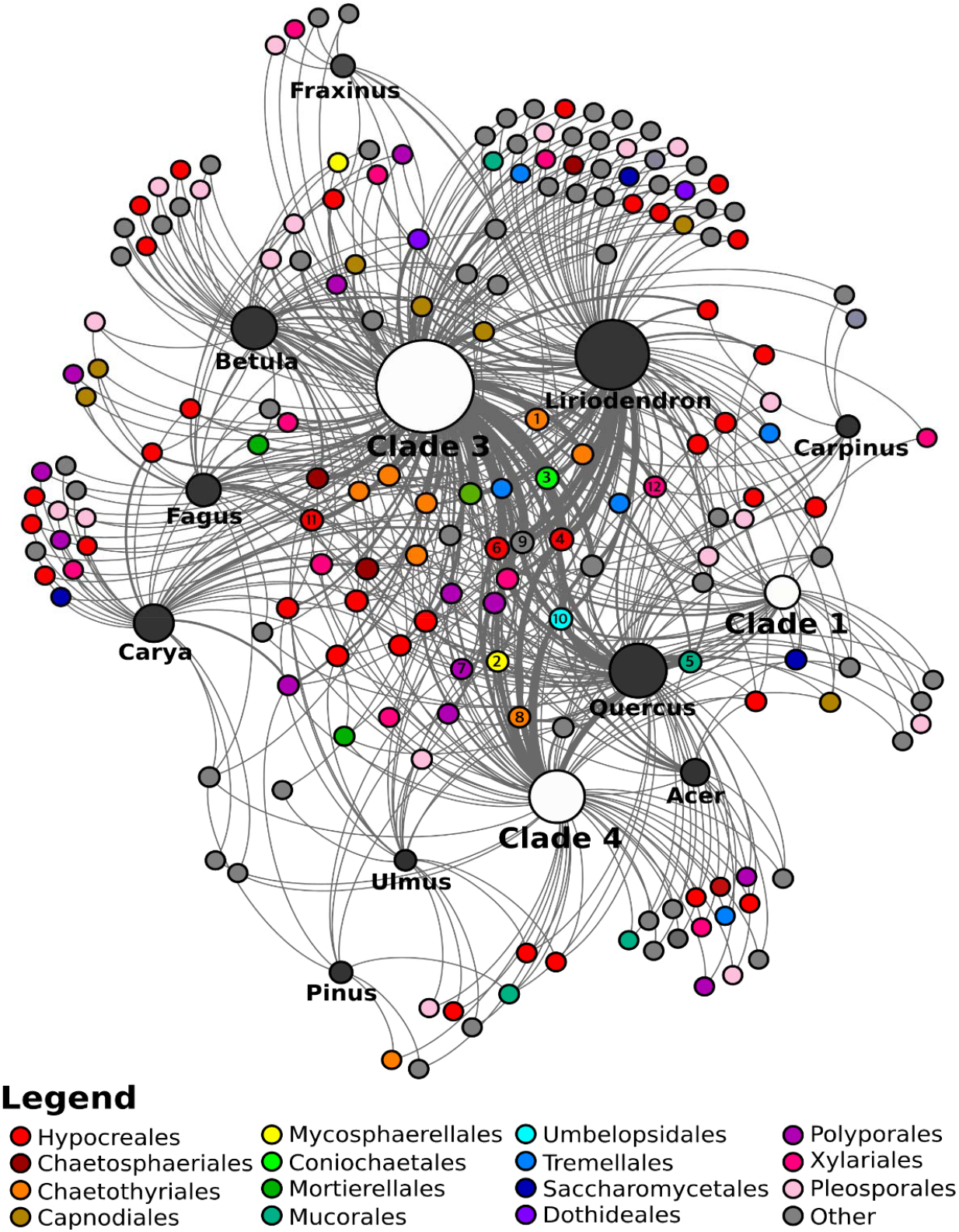
Fungal community network across *B. lecontii* clades and wood substrates. Small unlabeled nodes represent fungal genera, color-coded by taxonomic order. Orders with fewer than 10 isolates are lumped into “Other”. Genus nodes with betweenness-centrality scores >0.5 are labeled with the rank of their betweenness-centrality score (1-12). White and black nodes represent clades and wood substrates, respectively. For these, the size of the node represents the relative sample size. Edges represent interactions between a fungal genus and a wood substrate/clade. Edge boldness indicates the strength of the interaction.

The betweenness-centrality scores from the network revealed that the core of the Brachycybe-associated fungal community is comprised of a small group of fungal genera that is fairly representative of the diversity in the broader millipede-associated fungal community. The structure of the network indicates that these core fungi are consumed by many individuals from different lineages of *B. lecontii* across its reported range and across many wood substrates. As such, these fungi may be the preferred fungal food source for *B. lecontii*. Alternatively, these fungi may readily survive gut passage, which would result in them being over-represented after culturing.

Only a single member of the order Polyporales, *Phanerochaete,* falls in the core of the fungal network, a highly unexpected result given that the majority of community science records of *Brachycybe* interacting with fungi appear to show the millipedes associating with Polyporales and closely allied decay fungi (Supplemental Table 1). It is possible that these fungi may serve a vital role, despite their near-absence from the core network. One such role may be to precondition substrates for arthropod colonization. For example, vascular wilt fungi predispose trees to attack by wood-boring ambrosia beetles (Hulcr & Stelinski 2017). The Verticillium wilt pathogen, *V. nonalfalfae*, predisposes tree-of-heaven to mass colonization by the ambrosia beetle *Euwallacea validus.* However, the fungal community recovered from surface-disinfested beetles does not include *V. nonalfalfae* (Kasson *et al.* 2013). As such, *V. nonalfalfae* might be overlooked in its role to precondition substrates for arthropod colonization. A second possibility is that millipedes do actively utilize wood decay fungi but do not exhibit strict fidelity with single species or rely disproportionately on individual fungal community members (Kasson *et al.* 2013, Jusino *et al.* 2015, Jusino *et al.* 2016). Nevertheless, these results, much like studies examining fungal communities in red-cockaded woodpecker excavations, may indicate millipedes are either (1) selecting degraded logs with a pre-established preferred fungal community, or (2) selecting fresh logs without any evidence of decay, then subsequently facilitating colonization by specific fungi (Jusino *et al.* 2015, Jusino *et al.* 2016).

### 3.2. Pathogenicity testing

To determine how members of the Polyporales and other fungi interact with *B. lecontii,* millipedes were challenged with pure cultures of a representative set of fungal isolates for seven days. Only four of 21 isolates caused significant mortality after seven days (Table 2): *Metarhizium flavoviride* (Hypocreales; Log-rank p < 0.0005), *Bjerkandera adusta* (Polyporales; Log-rank p < 0.0005), *Irpex lacteus* (Polyporales; Log-rank p = 0.014), and *Trametopsis cervina* (Polyporales; Log-rank p < 0.0005). Interestingly, the known virulent entomopathogens *Lecanicillium attenuatum, Pochonia bulbillosa,* and *Verticillium insectorum* caused little to no mortality to *Brachycybe* after seven days of continuous exposure (Table 2).

Since three of the four pathogenic fungi were in the Polyporales, a follow-up experiment was performed using 18 isolates from the Polyporales, and one isolate of *Peniophora* (Russulales), the only fungus reported in the literature to be in association with *Brachycybe* (Gardner 1975). Only three of these isolates were significantly more pathogenic than the sterile agar control (Table 3): *Gloeoporus pannocinctus* (Polyporales; Log-rank p < 0.0007), and two isolates of *Trametopsis cervina* (Polyporales; Log-rank p for both < 0.0007). The *Bjerkandera* isolate and the *Irpex* isolate that were significantly pathogenic in the first assay were not significantly pathogenic in the second. However, the number of individuals used in the second assay was 10 per treatment, as compared to 15 per treatment in the first assay, and two individuals in the control treatment in the second assay were dead by the end of the assay. Together, these factors suggest that the results of the second assay are less reliable.

The results of the two pathogenicity assays indicate that several members of the Polyporales, including *Bjerkandera, Irpex, Trametopsis,* and *Gloeoporus,* may be pathogenic to *Brachycybe* millipedes (Figure 3), while three of the four entomopathogenic Hypocreales were not pathogenic. Additionally, seven other fungal orders caused little to no mortality (Table 2).

**Figure 3.**
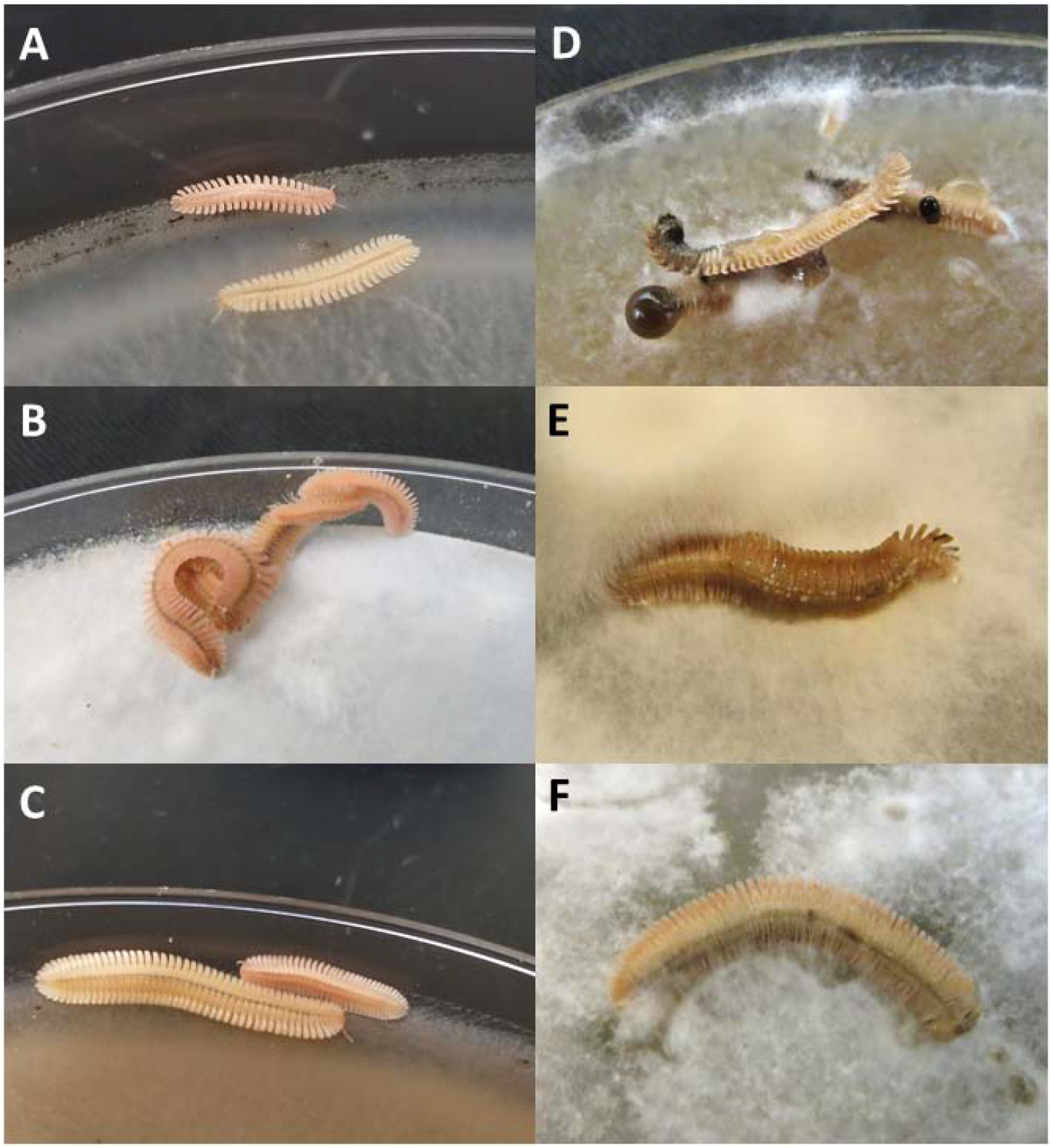
Representative outcomes of live-plating assay with Polyporales. A-C show *B. lecontii* with no outward disease symptoms after 7 days of exposure to the indicated fungus, and D-F show millipedes that were killed by the indicated fungus. In all three examples of mortality, fungal hyphae are growing over the millipede. A: *Phlebia livida* (BC0629), B: *Ceriporia lacerata* (BC1158), C: *Scopuloides rimosa* (BC1046), D: *Trametopsis cervina* (BC0494), E: *Bjerkandera adusta* (BC0310), F: *Irpex lacteus* (BC0523).

It is unclear how *B. lecontii* resists the well-documented entomopathogenic effects of many Hypocreales with the exception of *Metarhizium* (Hajek & St. Leger 1994). Parallel studies by Macias (2017) demonstrated that the Hypocrealean isolates used in the *Brachycybe* pathogenicity assays were pathogenic to insects. In contrast, the high incidence of pathogenicity among Brachycybe-associated Polyporales was unexpected. In a separate study, insects challenged with these same Polyporales were completely unaffected (Macias 2017).

One hypothesis is that Polyporales, depending on whether they are in a growth phase or a fruiting phase (Lu *et al.* 2014, Calvo *et al.* 2002), may produce chemicals that inadvertently harm millipedes. Since fruiting bodies were never observed in culture, it is likely that the fungi used in the pathogenicity assays were in a growth phase, which proved detrimental to *B. lecontii.* A second hypothesis that may explain pathogenicity among the Polyporales is that experiments relying on pure cultures of a single fungus do not account for fungus-fungus interactions or interactions between fungi and other organisms (Li & Zhang 2014, Macias 2017).

### 3.3. New species

At least seven putative new species were identified, but only two were investigated in this study. The five not examined are “aff. *Coniochaeta*” (Coniochaet ales), “aff. *Leptodontidium*” (Helotiales), “*Pseudonectria* aff. *buxi*” (Hypocreales), “aff. *Fonsecaea* sp.” (Chaetothyriales) and “aff. *Oidiodendron*” (Onygenales). The two examined species were from the phylum Mucoromycota (Spatafora *et al.* 2017), in the orders Mortierellales and Mucorales.

*Mortierella* aff. *ambigua* is represented by 27 clonal isolates from seven widespread collection sites (AR1, AR3, AR4, VA3, WV2, WV4, and WV5), five wood substrates (*Acer, Fagus, Fraxinus, Liriodendron,* and *Quercus*), and two millipede clades (Clade 3 and 4). These isolates are 92% identical to strain *“Mortierella ambigua* CBS 450.88” and were deposited as GenBank accessions MH971275 and MK045304 (Supplemental Table 2 & 4). All isolates of *“Mortierella* aff. *ambigua*” produced large gemmae (Aki *et al.* 2001, Embree 1963) as the cultures aged past 7 days. These structures grew to at most half a centimeter across and were present on the surface and embedded in the media (Figure 4). Sporangia were not observed in any of the *Mortierella* aff. *ambigua* isolates so comparisons with known *Mortierella* sporangial morphology could not be made. More in-depth morphological studies are needed before a formal description can be made.

**Figure 4.**
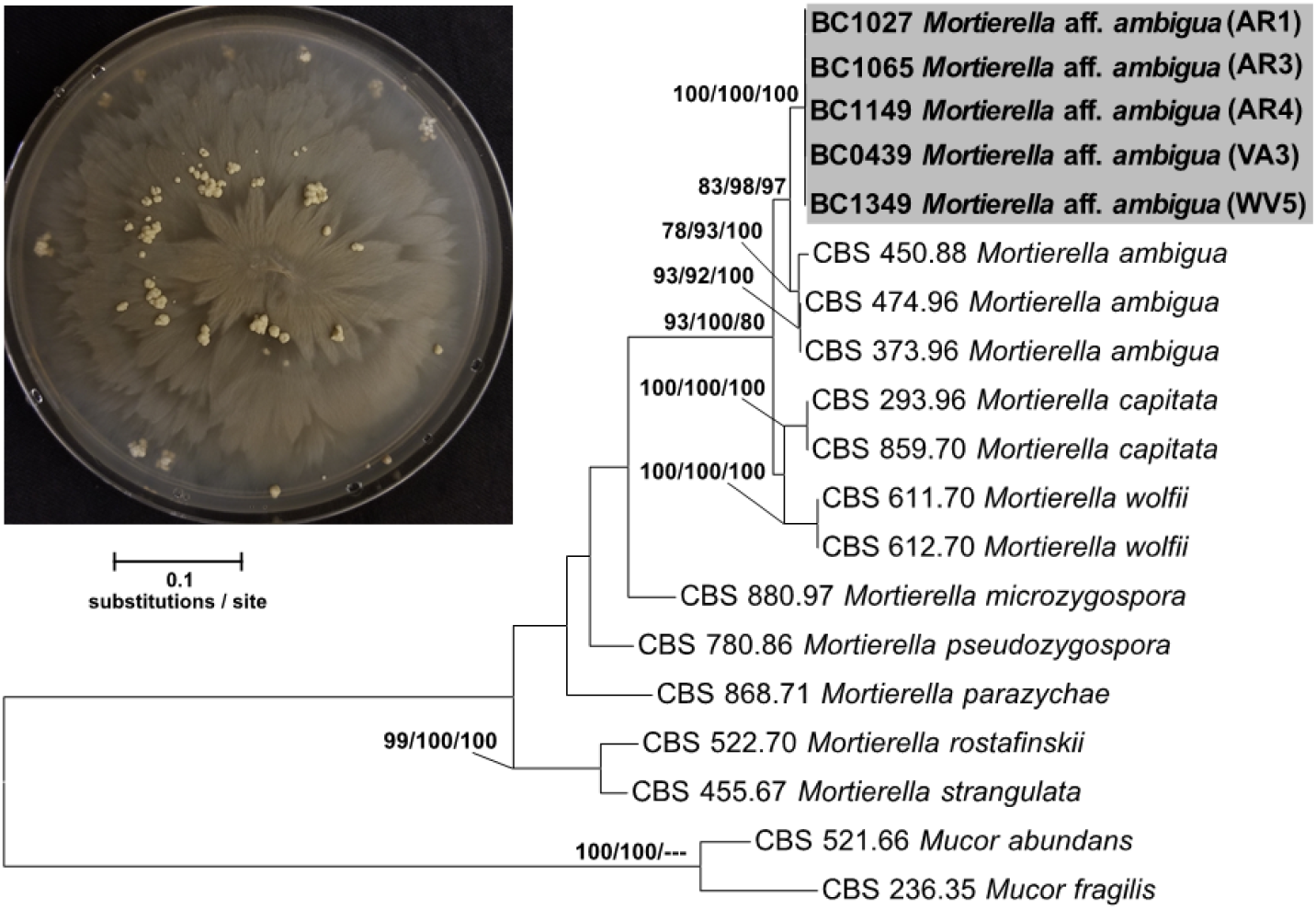
Concatenated ITS+LSU phylogenetic tree of *Mortierella* aff. *ambigua* and close relatives. Bootstrap support and posterior probabilities are indicated near each node (ML/MP/BI), and nodes with >50% support are labeled. Dashes indicate that a particular node did not appear in the indicated analysis. The grey box indicates the isolates belonging to *Mortierella* aff. *ambigua.* A representative culture of the fungus with gemmae is shown in the upper left.

The initial alignment included 1149 characters and the final dataset was reduced to 834 characters, and the maximum parsimony analysis yielded 7 most-parsimonious trees with a length of 732. Phylogenetic analysis of a concatenated ITS+LSU 17-isolate dataset including 5 *Mortierella* aff. *ambigua* confirmed placement of this novel species sister to *M. ambigua* sensu stricto (Figure 4) and inside the previously described Clade 5 of *Mortierella* (Wagner *et al.* 2013). Clade 5 *Mortierella* species are common from soil but have also been associated with amphipods and invasive mycoses in humans. More distantly related species such as *Mortierella beljakovae* and *Mortierella formicicola* have known associations with ants but the nature of this relationship remains unclear (Wagner *et al.* 2013).

The second putative new species, aff. *Apophysomyces* sp., is represented by five isolates from one site (OK1), one wood substrate *(Quercus),* and one millipede clade (Clade 4). These isolates are 84% identical to strain *“Apophysomyces ossiformis* strain UTHSC 04-838” and were deposited as GenBank accessions <u>MH971276</u> and <u>MK045305</u> (Supplemental Table 3 & 4). Sporangial morphology of these isolates aligns with described features for this genus (Alvarez *et al.* 2010, Bonifaz *et al.* 2014), but more in-depth morphological studies are needed.

The initial alignment included 1258 characters and the final dataset was reduced to 935 characters, and the maximum parsimony analysis yielded 3 most-parsimonious trees with a length of 1101. Phylogenetic analysis of a concatenated ITS+LSU 16 isolate dataset including three aff. *Apophysomyces* sp. isolates confirmed placement of this novel species sister to the clade containing known species of *Apophysomyces* sp. (Figure 5). The combined branch length among all known species of *Apophysomyces* (0.0589 substitutions/site) is less than the branch length separating our putative new species and these known species (0.0903 substitutions/site), providing evidence that our novel species is in fact, a novel genus. The genus *Apophysomyces* has been isolated from soil but is also known to cause severe mycoses in immunocompetent humans (Alvarez *et al.* 2010, Bonifaz *et al.* 2014, Etienne *et al.* 2012).

**Figure 5.**
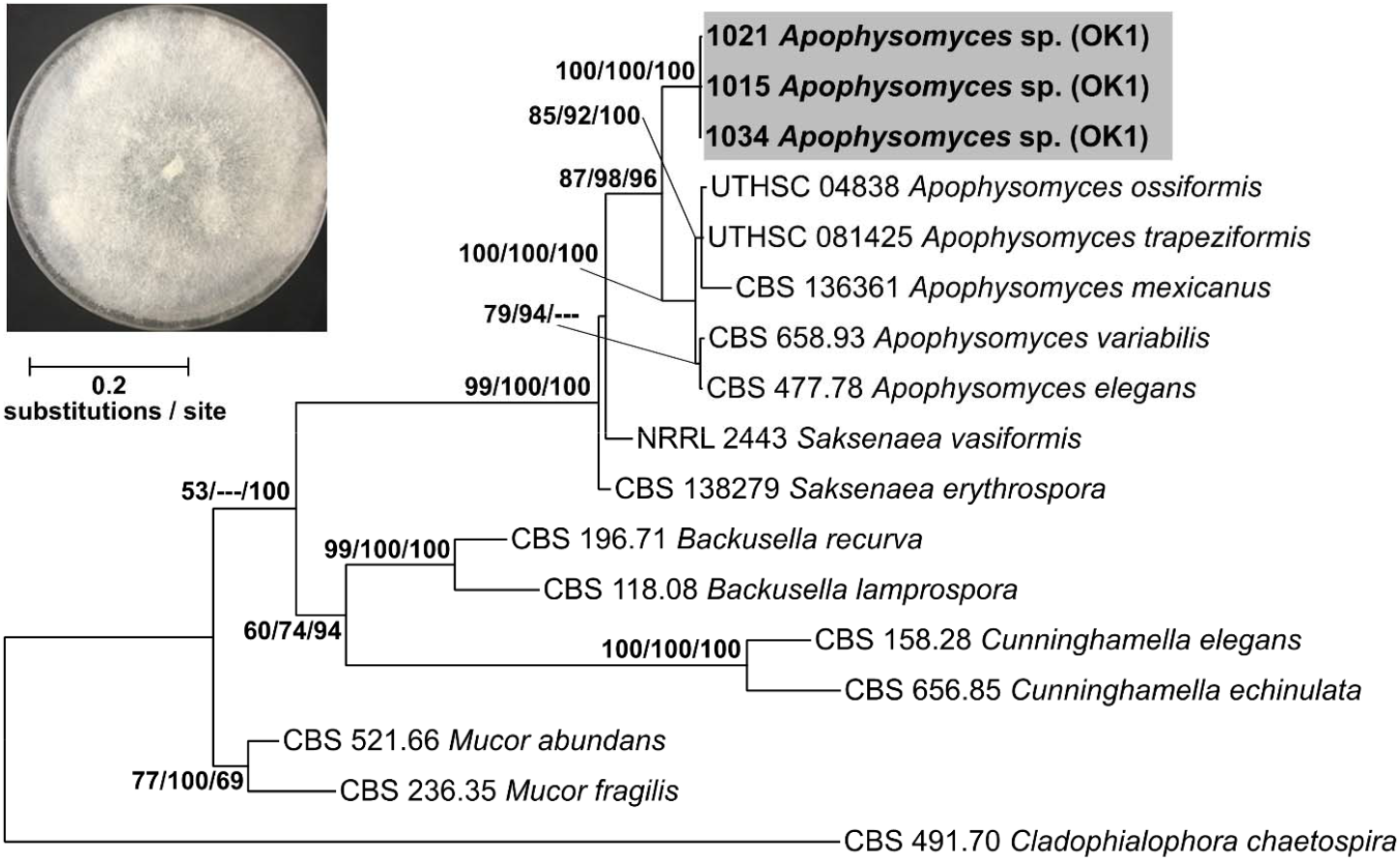
Concatenated ITS+LSU phylogenetic tree of aff. *Apophysomyces* sp. and close relatives. Bootstrap support and posterior probabilities are indicated near each node (ML/MP/BI), and nodes with >50% support are labeled. Dashes indicate that a particular node did not appear in the indicated analysis. The grey box indicates the isolates belonging to aff. *Apophysomyces* sp. A representative culture of the fungus is shown in the upper left.

Despite the use of more classical culture-based approaches, the recovery of seven putative new species highlights the vast amount of undescribed fungal biodiversity associated with millipedes. Culture-independent approaches will undoubtedly uncover many additional new species, possibly including some from unculturable lineages of fungi.

### 3.4. Conclusion

*Brachycybe lecontii* associates with a large and diverse community of fungi, including at least 176 genera in 39 fungal orders from four phyla. Significant differences in the fungal community among wood substrates, millipede clades, and ecoregions indicate that these factors influence the composition of millipede-associated fungal communities, while millipede sex does not. One putative new species and one putative new genus of fungi were found and examined in this study, and there is evidence for several additional new species that remain to be assessed phylogenetically. The core fungal community consists of fungi from at least nine orders, primarily members of phylum Ascomycota. While community science records of *Brachycybe* show the millipedes interacting with almost entirely Basidiomycota, especially Polyporales, only one genus from that order occurs in the core of the network. Four genera in the Polyporales were found to be pathogenic to *Brachycybe* in live-plating assays, while three genera of notorious entomopathogens from the Hypocreales did not cause significant mortality in millipedes. Only a single fungus outside the Polyporales caused significant mortality.

In less than a decade, the research on arthropod-fungus interactions has accelerated and led to the discoveries of several new associations (You *et al.* 2015, Menezes *et al.* 2015, Voglmayr *et al.* 2011). This study demonstrates that the complexity of millipede-fungus interactions has been underestimated and these interactions involve many undescribed species. This paper represents the first comprehensive survey of fungal communities associated with any member of the millipede subterclass Colobognatha. We anticipate that future studies of millipede-fungus interactions will yield countless new fungi and clarify the ecology these interactions.

## Acknowledgements

We thank Derek Hennen, Mark Double, Nicole Utano, and Toby Grapner for assistance with collection of millipedes and/or maintenance of cultured fungi. AM was supported by WVU Ruby Fellowship. MTK was supported by the West Virginia Agricultural and Forestry Experiment Station, Morgantown, WV.

## Author contributions

A.M.M., P.E.M., E.M.M., M.S.B., D.G.P., R.V.M.R., and M.T.K. conceived of the study. A.M.M., D.P.G.S., C.M.S., K.L.W., M.C.B., A. M. M., V.W., T.H.J., and M.T.K. performed laboratory work with the help/advice of P.E.M., M.C.B., D.G.P., J.E.S., G.R.B. A.M.M., P.E.M., E.M.M., and M.T.K. analyzed data. A.M.M., P.E.M., E.M.M., M.S.B., J.E.S., G.R.B, R.V.M.R., and M.T.K. wrote the manuscript with input from all coauthors.

## Supplementary data

Supplementary data related to this article can be found at (enter link here).

