## Supplemental Tables 1-4 for "Diversity and function of fungi associated with the fungivorous millipede, *Brachycybe lecontii*"

Supplemental Table 1. Community science records of *Brachycybe* from BugGuide and iNaturalist websites. Habit and fungus columns were based on subjective evaluations of associated images.

| Species | Database | URL | Year | Contributor | Habit | Fungus? |
| --- | --- | --- | --- | --- | --- | --- |
| *B. lecontii* | BugGuide | https://bugguide.net/node/view/45/bgpage | 2002 | Troy Bartlett | Colony on bark | Not apparent |
| *B. lecontii* | BugGuide | https://bugguide.net/node/view/49/bgpage | 2000 | Troy Bartlett | Colony on bark | Not apparent |
| *B. lecontii* | BugGuide | https://bugguide.net/node/view/57554/bgpage | 1969 | Steven Barney | Individual on dead leaf | Not apparent |
| *B. lecontii* | BugGuide | https://bugguide.net/node/view/213351/bgpage | 2008 | Tim Nichols | Colony on bark (?) | Not apparent |
| *B. lecontii* | BugGuide | https://bugguide.net/node/view/595215/bgpage | 2011 | David J. Thomas | Individual on rock | Not apparent |
| *B. lecontii* | BugGuide | https://bugguide.net/node/view/1178937/bgpage | 2016 | Jonathan Carpenter | Individual on dead wood | Not apparent |
| *B. lecontii* | BugGuide | https://bugguide.net/node/view/1517909/bgpage | 2018 | John Lampkin | Individual on rock | Not apparent |
| *B. lecontii* | iNaturalist | https://www.inaturalist.org/observations/17512358 | 2018 | "wjneely" | (Individual in collection) | Not apparent |
| *B. lecontii* | iNaturalist | https://www.inaturalist.org/observations/17283380 | 2018 | "rhinoclemmys" | Individual on dead wood | Not apparent |
| *B. lecontii* | iNaturalist | https://www.inaturalist.org/observations/13439380 | 2018 | "kevinfitzpatrick" | (Individual in collection) | Not apparent |
| *B. lecontii* | iNaturalist | https://www.inaturalist.org/observations/12280088 | 2018 | "cavemander17 " | Individual on rock | Not apparent |
| *B. lecontii* | iNaturalist | https://www.inaturalist.org/observations/10718361 | 2018 | "paulmarek" | Individual on dead wood | Not apparent |
| *B. lecontii* | iNaturalist | https://www.inaturalist.org/observations/8057119 | 2017 | "hazelsnail" | (Individual in collection) | Not apparent |
| *B. lecontii* | iNaturalist | https://www.inaturalist.org/observations/6737379 | 2017 | "beschwar" | Colony on rotten wood | Not apparent |
| *B. lecontii* | iNaturalist | https://www.inaturalist.org/observations/6604286 | 2017 | "mark_swanson" | Individual on dead wood | Not apparent |
| *B. lecontii* | iNaturalist | https://www.inaturalist.org/observations/6537354 | 2017 | "libbing_life" | Individual on dead wood | Not apparent |
| *B. lecontii* | iNaturalist | https://www.inaturalist.org/observations/6451167 | 2017 | "reallifeecology" | Individual on dead wood | Not apparent |
| *B. lecontii* | iNaturalist | https://www.inaturalist.org/observations/6451129 | 2017 | "muir" | Individual on dead wood | Not apparent |
| *B. lecontii* | iNaturalist | https://www.inaturalist.org/observations/5524007 | 2017 | "chinquapin" | Individual on wood | Not apparent |
| *B. lecontii* | iNaturalist | https://www.inaturalist.org/observations/5012087 | 2017 | "reallifeecology" | Individual on dead wood | Not apparent |
| *B. lecontii* | iNaturalist | https://www.inaturalist.org/observations/4534657 | 2016 | "reallifeecology" | Individual on rock | Not apparent |
| *B. lecontii* | iNaturalist | https://www.inaturalist.org/observations/3402635 | 2016 | "mhedin" | Colony on rotten wood | Not apparent |
| *B. lecontii* | iNaturalist | https://www.inaturalist.org/observations/3007821 | 2016 | "reallifeecology" | (Individual in collection) | Not apparent |
| *B. lecontii* | iNaturalist | https://www.inaturalist.org/observations/2877858 | 2016 | "damontighe" | Large colony on dead wood | Not apparent |
| *B. lecontii* | iNaturalist | https://www.inaturalist.org/observations/453193 | 2013 | "eric_hunt" | Individual on dead wood | Not apparent |
| *B. lecontii* | iNaturalist | https://www.inaturalist.org/observations/134815 | 2009 | "rcurtis" | (Individual in collection) | Not apparent |
| *B. lecontii* | iNaturalist | https://www.inaturalist.org/observations/4308248 | 2014 | "michaelskvarla" | Colony on dead wood | Yes; grey phlebioid crust |
| *B. lecontii* | iNaturalist | https://www.inaturalist.org/observations/11292069 | 2018 | "teriyaki12" | Colony on rotten wood | Yes; tan poroid crust |
| *B. lecontii* | iNaturalist | https://www.inaturalist.org/observations/2360344 | 2015 | "eric_hunt" | Colony on dead wood | Yes; *Terana caerulea* |
| *B. lecontii* | iNaturalist | https://www.inaturalist.org/observations/13127260 | 2018 | "rdandekar" | Colony on dead wood | Yes; *Trichoderma*-infected white phlebioid crust |
| *B. lecontii* | iNaturalist | https://www.inaturalist.org/observations/11265140 | 2018 | "goody" | Colony on dead wood | Yes; well-rotted black fungus |
| *B. lecontii* | iNaturalist | https://www.inaturalist.org/observations/12606811 | 2018 | "jeremysouthers1" | Individual on pine bark | Yes; white crust |
| *B. lecontii* | iNaturalist | https://www.inaturalist.org/observations/9988932 | 2017 | "rdandekar" | Colony on dead wood | Yes; white crust |
| *B. lecontii* | iNaturalist | https://www.inaturalist.org/observations/5322157 | 2005 | "larry14" | Colony on dead wood | Yes; white crust |
| *B. lecontii* | iNaturalist | https://www.inaturalist.org/observations/5212886 | 2017 | "muddynaturalist" | Colony on dead wood | Yes; white hyphal cords |
| *B. lecontii* | iNaturalist | https://www.inaturalist.org/observations/3915382 | 2016 | "chris184" | Colony on dead wood | Yes; white hyphal mat and cup fungus |
| *B. lecontii* | iNaturalist | https://www.inaturalist.org/observations/13292468 | 2018 | "daniel_folds " | Large colony on snag | Yes; white poroid bracket |
| *B. lecontii* | BugGuide | https://bugguide.net/node/view/53529/bgimage | 2006 | Jonathan Burishkin | Colony on dead wood | Yes; white-grey crust |
| *B. petasata* | BugGuide | https://bugguide.net/node/view/270869/bgimage | 2009 | Rob Craig | Colony under rotten log | Yes; phlebioid white crust |
| *B. producta* | BugGuide | https://bugguide.net/node/view/1027823/bgimage | 2014 | Sam McNally | Individual on dead wood | Not apparent |
| *B. producta* | iNaturalist | https://www.inaturalist.org/observations/10618906 | 2018 | "damarisb" | Individual on wood | Not apparent |
| *B. producta* | iNaturalist | https://www.inaturalist.org/observations/9480009 | 2018 | "kueda" | Individual on dead wood | Not apparent |
| *B. producta* | iNaturalist | https://www.inaturalist.org/observations/9465211 | 2018 | "tiwane" | Individual on dead wood | Not apparent |
| *B. producta* | iNaturalist | https://www.inaturalist.org/observations/8771365 | 2017 | "mazer" | Colony on rotten wood | Not apparent |
| *B. producta* | iNaturalist | https://www.inaturalist.org/observations/5540144 | 2017 | "richardwasson" | (Individual in collection) | Not apparent |
| *B. producta* | iNaturalist | https://www.inaturalist.org/observations/5536177 | 2017 | "dominic" | Colony on rotten wood | Not apparent |
| *B. producta* | iNaturalist | https://www.inaturalist.org/observations/4502473 | 2016 | "icosahedron" | Colony on dead wood | Not apparent |
| *B. producta* | iNaturalist | https://www.inaturalist.org/observations/1404122 | 2015 | "loarie" | Individual on dead wood | Not apparent |
| *B. producta* | iNaturalist | https://www.inaturalist.org/observations/1117299 | 2014 | "biosam" | Individual on dead wood | Not apparent |
| *B. producta* | iNaturalist | https://www.inaturalist.org/observations/1108594 | 2011 | "temminicki" | Individual on dead wood | Not apparent |
| *B. producta* | BugGuide | https://bugguide.net/node/view/514775/bgimage | 2011 | Timothy Boomer | Colony on rotten wood | Yes; brown crust |
| *B. producta* | iNaturalist | https://www.inaturalist.org/observations/4551536 | 2016 | "m_patton" | Colony on dead wood | Yes; brown crust |
| *B. producta* | iNaturalist | https://www.inaturalist.org/observations/5406781 | 2017 | "catchang" | Colony on rotten wood | Yes; grey hyphae |
| *B. producta* | iNaturalist | https://www.inaturalist.org/observations/5471277 | 2017 | "loarie" | Colony on rotten wood | Yes; tan phlebioid crust |
| *B. producta* | iNaturalist | https://www.inaturalist.org/observations/10189738 | 2018 | "katewread" | Colony on rotten wood | Yes; white crust |
| *B. producta* | iNaturalist | https://www.inaturalist.org/observations/9556670 | 2018 | "lisahug" | Individual on dead wood | Yes; white crust |
| *B. producta* | iNaturalist | https://www.inaturalist.org/observations/2701345 | 2016 | "loarie" | Individual on dead wood | Yes; white hyphal mat |
| *B. producta* | iNaturalist | https://www.inaturalist.org/observations/6908225 | 2017 | "biosam" | Individual on rotten wood | Yes; white-green hyphal mat |
| *B. producta* | iNaturalist | https://www.inaturalist.org/observations/4784269 | 2016 | "icosahedron" | Colony on dead wood | Yes; white-tan crust |
| *B. rosea* | BugGuide | https://bugguide.net/node/view/169590/bgpage | 2007 | Jim McClarin | Rotting oak wood | Not apparent |
| *B. rosea* | BugGuide | https://bugguide.net/node/view/418663/bgpage | 2010 | Debbi Brusco | Individual on dead wood | Not apparent |
| *B. rosea* | BugGuide | https://bugguide.net/node/view/1508463/bgimage | 2018 | "Madratter" | Individual on dead leaf | Not apparent |
| *B. rosea* | BugGuide | https://bugguide.net/node/view/904123/bgimage | 2014 | Sharon Hadley | Individual on dead wood | Not apparent |
| *B. rosea* | iNaturalist | https://www.inaturalist.org/observations/6543298 | 2017 | "biosam" | (Colony in collection) | Not apparent |
| *B. rosea* | iNaturalist | https://www.inaturalist.org/observations/5914163 | 2017 | "leptonia" | Individual on dead wood | Not apparent |
| *B. rosea* | iNaturalist | https://www.inaturalist.org/observations/3067933 | 2016 | "rebeccafay" | Colony on rotten wood | Not apparent |
| *B. rosea* | iNaturalist | https://www.inaturalist.org/observations/2896872 | 2016 | "mhedin" | Individual on wood | Not apparent |
| *B. rosea* | iNaturalist | https://www.inaturalist.org/observations/1310859 | 2015 | "kueda" | Colony on dead wood | Not apparent |
| *B. rosea* | iNaturalist | https://www.inaturalist.org/observations/1205992 | 2015 | "kueda" | Individual on dead wood | Not apparent |
| *B. rosea* | iNaturalist | https://www.inaturalist.org/observations/23559 | 2011 | "biosam" | Colony on dead wood | Not apparent |
| *B. rosea* | iNaturalist | https://www.inaturalist.org/observations/4039035 | 2016 | "damontighe" | Individual on dead wood | Yes; white hyphal mat |
| *B.* sp. | BugGuide | https://bugguide.net/node/view/1459279/bgimage | 2017 | Bob Kipfer | Individual on dead wood | Not apparent |
| *B.* sp. | BugGuide | https://bugguide.net/node/view/1421004/bgimage | 2017 | "Hobo Joe A.K.A. Insect Lover" | Individual on dead wood | Not apparent |
| *B.* sp. | BugGuide | https://bugguide.net/node/view/1104425/bgimage | 2015 | Jonathan Carpenter | (Individual in collection) | Not apparent |
| *B.* sp. | BugGuide | https://bugguide.net/node/view/865459/bgimage | 2013 | Mark H Brown | Individual on rock | Not apparent |
| *B.* sp. | BugGuide | https://bugguide.net/node/view/843152/bgimage | 2013 | William Hull | Individual on dead wood | Not apparent |
| *B.* sp. | BugGuide | https://bugguide.net/node/view/784265/bgimage | 2013 | "BugCatcher10" | (Individual in collection) | Not apparent |
| *B.* sp. | BugGuide | https://bugguide.net/node/view/620056/bgimage | 2012 | Marvin Smith | Colony on soil | Not apparent |
| *B.* sp. | BugGuide | https://bugguide.net/node/view/413806/bgimage | 2010 | Scott Cox | Individual on wood | Not apparent |
| *B.* sp. | BugGuide | https://bugguide.net/node/view/176887/bgimage | 2008 | Natalie McNear | Individual on dead wood | Not apparent |
| *B.* sp. | BugGuide | https://bugguide.net/node/view/172005/bgimage | 2008 | Natalie McNear | Colony on rotten wood | Not apparent |
| *B.* sp. | iNaturalist | https://www.inaturalist.org/observations/17666548 | 2018 | "sageharmon" | (Individual in collection) | Not apparent |
| *B.* sp. | iNaturalist | https://www.inaturalist.org/observations/17422879 | 2018 | "ecology2" | (Individual in collection) | Not apparent |
| *B.* sp. | iNaturalist | https://www.inaturalist.org/observations/17418303 | 2018 | "jainitastic" | (Individual in collection) | Not apparent |
| *B.* sp. | iNaturalist | https://www.inaturalist.org/observations/12322132 | 2018 | "apteryxrowi" | (Individual in collection) | Not apparent |
| *B.* sp. | iNaturalist | https://www.inaturalist.org/observations/12152016 | 2018 | "lorri-gong" | Individual on dead wood | Not apparent |
| *B.* sp. | iNaturalist | https://www.inaturalist.org/observations/11718802 | 2018 | "magicicada" | Individual on dead wood | Not apparent |
| *B.* sp. | iNaturalist | https://www.inaturalist.org/observations/11510349 | 2018 | "kestrel" | Individual on dead wood | Not apparent |
| *B.* sp. | iNaturalist | https://www.inaturalist.org/observations/11095607 | 2018 | "garmonb0zia" | Individual on dead wood | Not apparent |
| *B.* sp. | iNaturalist | https://www.inaturalist.org/observations/10940260 | 2018 | "christopher13" | Colony on rotten wood | Not apparent |
| *B.* sp. | iNaturalist | https://www.inaturalist.org/observations/10178482 | 2018 | "christopher13" | Individual on wood | Not apparent |
| *B.* sp. | iNaturalist | https://www.inaturalist.org/observations/9963155 | 2018 | "l0000_hz_legend" | Colony on rotten wood | Not apparent |
| *B.* sp. | iNaturalist | https://www.inaturalist.org/observations/9307728 | 2017 | "irislane" | Individual on dead wood | Not apparent |
| *B.* sp. | iNaturalist | https://www.inaturalist.org/observations/9307713 | 2017 | "irislane" | Individual on dead wood | Not apparent |
| *B.* sp. | iNaturalist | https://www.inaturalist.org/observations/9281269 | 2017 | "temminicki" | Individual on dead wood | Not apparent |
| *B.* sp. | iNaturalist | https://www.inaturalist.org/observations/8993568 | 2017 | "easmeds" | Colony on bark | Not apparent |
| *B.* sp. | iNaturalist | https://www.inaturalist.org/observations/7609652 | 2017 | "tonytravlos" | Colony on rotten wood | Not apparent |
| *B.* sp. | iNaturalist | https://www.inaturalist.org/observations/6937088 | 2017 | "biosam" | Colony on rotten wood | Not apparent |
| *B.* sp. | iNaturalist | https://www.inaturalist.org/observations/6253831 | 2017 | "chinquapin" | Colony on soil | Not apparent |
| *B.* sp. | iNaturalist | https://www.inaturalist.org/observations/6207443 | 2017 | "friel" | Individual on rotten wood | Not apparent |
| *B.* sp. | iNaturalist | https://www.inaturalist.org/observations/5722971 | 2017 | "geodani" | (Individual in collection) | Not apparent |
| *B.* sp. | iNaturalist | https://www.inaturalist.org/observations/5714347 | 2017 | "lorri-gong" | Individual on dead wood | Not apparent |
| *B.* sp. | iNaturalist | https://www.inaturalist.org/observations/5534690 | 2017 | "christopher13" | Colony on rotten wood | Not apparent |
| *B.* sp. | iNaturalist | https://www.inaturalist.org/observations/5282983 | 2016 | "sclerobunus" | Individual on dead wood | Not apparent |
| *B.* sp. | iNaturalist | https://www.inaturalist.org/observations/5228142 | 2017 | "vermfly" | Individual on dead wood | Not apparent |
| *B.* sp. | iNaturalist | https://www.inaturalist.org/observations/5189159 | 2017 | "christopher13" | Individual on dead wood | Not apparent |
| *B.* sp. | iNaturalist | https://www.inaturalist.org/observations/2936552 | 2016 | "allisonpeters" | Individual on rotten wood | Not apparent |
| *B.* sp. | iNaturalist | https://www.inaturalist.org/observations/2730609 | 2016 | "marionanoiram" | Individual on dead wood | Not apparent |
| *B.* sp. | iNaturalist | https://www.inaturalist.org/observations/1522694 | 2015 | "beeboy" | Colony on dead wood | Not apparent |
| *B.* sp. | iNaturalist | https://www.inaturalist.org/observations/1333933 | 2015 | "damontighe" | Individual on dead leaf | Not apparent |
| *B.* sp. | iNaturalist | https://www.inaturalist.org/observations/158981 | 2007 | "kucycads" | Individual on dead wood | Not apparent |
| *B.* sp. | iNaturalist | https://www.inaturalist.org/observations/18981 | 2010 | "tapbirds" | Individual on dead wood | Not apparent |
| *B.* sp. | iNaturalist | https://www.inaturalist.org/observations/10674619 | 2006 | "henkwallays2" | Individual on wood | Yes; black-orange crust |
| *B.* sp. | iNaturalist | https://www.inaturalist.org/observations/1275824 | 2015 | "robberfly" | Colony on dead wood | Yes; cup fungus |
| *B.* sp. | iNaturalist | https://www.inaturalist.org/observations/9013075 | 2017 | "easmeds" | Individual on dead wood | Yes; gelatinous grey crust |
| *B.* sp. | iNaturalist | https://www.inaturalist.org/observations/14057280 | 2018 | "clinchriverdreams" | Individual on dead wood | Yes; grey crust |
| *B.* sp. | iNaturalist | https://www.inaturalist.org/observations/363144 | 2013 | "tonyg" | Colony on dead wood | Yes; grey crust |
| *B.* sp. | iNaturalist | https://www.inaturalist.org/observations/9722536 | 2018 | "kookamongus" | Individual on dead wood | Yes; grey phlebioid crust |
| *B.* sp. | iNaturalist | https://www.inaturalist.org/observations/5823516 | 2017 | "leenash" | Individual on dead wood | Yes; large brown crust |
| *B.* sp. | iNaturalist | https://www.inaturalist.org/observations/12265670 | 2018 | "athensalive" | Colony on rotten wood | Yes; *Phlebia* sp. |
| *B.* sp. | iNaturalist | https://www.inaturalist.org/observations/595969 | 2014 | "moonlittrails" | Colony on dead wood | Yes; phlebioid crust and cup fungus |
| *B.* sp. | iNaturalist | https://www.inaturalist.org/observations/5408733 | 2017 | "damontighe" | Individual on dead wood | Yes; phlebioid grey crust and white hyphal mat |
| *B.* sp. | iNaturalist | https://www.inaturalist.org/observations/11719631 | 2018 | "magicicada" | Individual on dead wood | Yes; pink-white hyphal mat |
| *B.* sp. | iNaturalist | https://www.inaturalist.org/observations/8870068 | 2017 | "debk" | Individual on rotten wood | Yes; tan bracket fungus and white poroid crust |
| *B.* sp. | iNaturalist | https://www.inaturalist.org/observations/12367836 | 2018 | "dgreenberger" | Colony on rotten wood | Yes; tan crust |
| *B.* sp. | iNaturalist | https://www.inaturalist.org/observations/5257850 | 2016 | "eccentric_entomophile" | Individual on dead wood | Yes; tan crust |
| *B.* sp. | iNaturalist | https://www.inaturalist.org/observations/10698727 | 2006 | "henkwallays2" | Large colony on dead wood | Yes; tan poroid crust |
| *B.* sp. | iNaturalist | https://www.inaturalist.org/observations/5452255 | 2017 | "damontighe" | Individual on dead wood | Yes; very rotten yellow crust |
| *B.* sp. | BugGuide | https://bugguide.net/node/view/851644/bgimage | 2011 | Sam McNally | Large colony on dead wood | Yes; white crust |
| *B.* sp. | iNaturalist | https://www.inaturalist.org/observations/9736972 | 2018 | "twillrichardson" | Colony on pine log | Yes; white crust |
| *B.* sp. | iNaturalist | https://www.inaturalist.org/observations/9281266 | 2017 | "temminicki" | Individual on dead wood | Yes; white crust |
| *B.* sp. | iNaturalist | https://www.inaturalist.org/observations/5920644 | 2017 | "leslie_flint" | Colony under rotten log | Yes; white crust |
| *B.* sp. | iNaturalist | https://www.inaturalist.org/observations/5229632 | 2017 | "tomv" | Colony on dead wood | Yes; white crust |
| *B.* sp. | iNaturalist | https://www.inaturalist.org/observations/2801341 | 2016 | "robberfly" | Colony on rotten wood | Yes; white crust |
| *B.* sp. | iNaturalist | https://www.inaturalist.org/observations/5095109 | 2016 | "mazer" | Colony on dead wood | Yes; white crust and phlebioid grey crust |
| *B.* sp. | iNaturalist | https://www.inaturalist.org/observations/9964190 | 2018 | "l0000_hz_legend" | Large colony on dead wood | Yes; white hyphae and orange crust |
| *B.* sp. | iNaturalist | https://www.inaturalist.org/observations/5408600 | 2017 | "robberfly" | Colony on rotten wood | Yes; white hyphal cords |
| *B.* sp. | iNaturalist | https://www.inaturalist.org/observations/3520995 | 2016 | "biosam" | Colony on rotten wood | Yes; white hyphal cords |
| *B.* sp. | iNaturalist | https://www.inaturalist.org/observations/3198564 | 2014 | "eccentric_entomophile" | Colony on rotten wood | Yes; white hyphal cords |
| *B.* sp. | iNaturalist | https://www.inaturalist.org/observations/1110059 | 2014 | "lindynik" | Colony on dead wood | Yes; white hyphal cords |
| *B.* sp. | iNaturalist | https://www.inaturalist.org/observations/1066156 | 2014 | "bapeck8" | Individual on dead wood | Yes; white hyphal cords |
| *B.* sp. | iNaturalist | https://www.inaturalist.org/observations/9812105 | 2018 | "amacedo" | Colony on rotten wood | Yes; white hyphal mat |
| *B.* sp. | iNaturalist | https://www.inaturalist.org/observations/8858584 | 2008 | "robirwin" | Individual on dead wood | Yes; white hyphal mat |
| *B.* sp. | iNaturalist | https://www.inaturalist.org/observations/7489413 | 2008 | "robirwin" | Individual on dead wood | Yes; white hyphal mat |
| *B.* sp. | iNaturalist | https://www.inaturalist.org/observations/6682063 | 2017 | "biosam" | Colony on rotten wood | Yes; white hyphal mat |
| *B.* sp. | iNaturalist | https://www.inaturalist.org/observations/5278798 | 2017 | "jessefurrow" | Individual on dead wood | Yes; white hyphal mat |
| *B.* sp. | iNaturalist | https://www.inaturalist.org/observations/5225939 | 2017 | "lenaz" | Individual on dead wood | Yes; white hyphal mat |
| *B.* sp. | iNaturalist | https://www.inaturalist.org/observations/4780656 | 2016 | "lorri-gong" | Colony on dead wood | Yes; white hyphal mat |
| *B.* sp. | iNaturalist | https://www.inaturalist.org/observations/1275884 | 2015 | "metsa" | Colony on dead wood | Yes; white hyphal mat |
| *B.* sp. | iNaturalist | https://www.inaturalist.org/observations/4505555 | 2016 | "dutchflatterry" | Colony on dead wood | Yes; white hyphal mat, tan bracket, slime mold |
| *B.* sp. | BugGuide | https://bugguide.net/node/view/945769/bgimage | 2014 | Jonathan Carpenter | Colony on rotten wood | Yes; white poroid crust |
| *B.* sp. | BugGuide | https://bugguide.net/node/view/633418/bgimage | 2012 | Alan Rockefeller | Colony on rotten wood | Yes; *Xylodon* sp. |
| *B.* sp. | iNaturalist | https://www.inaturalist.org/observations/5786153 | 2017 | "daniam" | Individual on dead wood | Yes; yellow slime mold and white hyphal cords |
| *B.* sp. | iNaturalist | https://www.inaturalist.org/observations/4902929 | 2017 | "grayson" | Colony on dead pine wood | Yes; yellow slime mold or cup fungus |

Supplemental Table 2. Strains and their associated NCBI Genbank reference numbers for isolates of *Mortierella* aff. *ambigua* and close relatives used in phylogenetic analysis.

| Name | Strain | ITS | LSU |
| --- | --- | --- | --- |
| *Mortierella ambigua* | CBS 450.88 | JX976067 | KC018411 |
| *Mortierella ambigua* | CBS 474.96 | JX976056 | KC018416 |
| *Mortierella ambigua* | CBS 373.96 | JX976062 | JX976147 |
| *Mortierella capitata* | CBS 293.96 | JX976123 | KC018334 |
| *Mortierella capitata* | CBS 859.70 | JX976008 | KC018395 |
| *Mortierella microzygospora* | CBS 880.97 | NR_111569 | HQ667394 |
| *Mortierella parazychae* | CBS 868.71 | HQ630283 | HQ667362 |
| *Mortierella pseudozygospora* | CBS 780.86 | JX975880 | JX976143 |
| *Mortierella wolfii* | CBS 611.70 | JN943806 | JN940863 |
| *Mortierella wolfii* | CBS 612.70 | MH859876 | HQ667381 |
| *Mortierella rostafinskii* | CBS 522.70 | NR_111586 | NG_042570 |
| *Mortierella strangulata* | CBS 455.67 | HQ630359 | HQ667437 |
| *Mucor abundans* | CBS 521.66 | JN206110 | JN206457 |
| *Mucor fragilis* | CBS 236.35 | JN205979 | FN650671 |

Supplemental Table 3. Strains and their associated NCBI Genbank reference numbers for isolates of aff. *Apophysomyces* sp. and close relatives used in phylogenetic analysis.

| Name | Strain number | ITS | LSU |
| --- | --- | --- | --- |
| *Apophysomyces variabilis* | CBS 658.93 | NR_130683 | HM849695 |
| *Apophysomyces elegans* | CBS 477.78 | JN206280 | JN206536 |
| *Apophysomyces ossiformis* | UTHSC 04-838 | NR_137035 | FN554252 |
| *Apophysomyces trapeziformis* | UTHSC 08-1425 | NR_137034 | FN554261 |
| *Apophysomyces mexicanus* | CBS 136361 | HG974255 | HG974256 |
| *Saksenaea vasiformis* | NRRL 2443 | FR687327 | HM776679 |
| *Saksenaea erythrospora* | CBS 138279 | KM102733 | KM102734 |
| *Mucor abundans* | CBS 521.66 | JN206110 | JN206457 |
| *Mucor fragilis* | CBS 236.35 | JN205979 | FN650671 |
| *Backusella circina* | CBS 128.70 | NR_103649 | JN206529 |
| *Backusella recurva* | CBS 196.71 | JN206265 | JN206523 |
| *Backusella lamprospora* | CBS 118.08 | NR_145291 | JN206531 |
| *Cunninghamella echinulata* | CBS 656.85 | JN205896 | JN206598 |
| *Cunninghamella elegans* | CBS 158.28 | JN205888 | JN206602 |
| *Cunninghamella bertholletiae* | CBS 190.84 | JN205878 | HM849701 |
| *Cladophialophora chaetospira* | CBS 491.70 | EU035405 | EU035405 |

Supplemental Table 4. Fungi recovered from *B. lecontii* based on ITS barcoding and arranged taxonomically.

| Name | |  |  | Example isolate | Best match | Query coverage (%) | Identity (%) | Deposited ITS sequence # |
| --- | --- | --- | --- | --- | --- | --- | --- | --- |
| Ascomycota | | |  |  |  |  |  |  |
|  | Amphisphaeriales | |  |  |  |  |  |  |
|  |  | *Discosia* | sp. | ---------- | --------------- | ---- | ---- | -------------- |
|  |  | *Neopestalotiopsis* | sp. | ---------- | --------------- | ---- | ---- | -------------- |
|  |  | *Pestalotiopsis* | *crassiuscula* | BC1288 | AY687868.1 | 100 | 99 | -------------- |
|  |  |  | *jesteri* | BC1501 | KT000165.1 | 95 | 99 | -------------- |
|  |  |  | *knightiae* | BC0084 | KM199311.1 | 99 | 99 | -------------- |
|  |  |  | *mangiferae* | BC1427 | KP074973.1 | 100 | 100 | -------------- |
|  |  |  | *microspora* | BC1439 | MH707065.1 | 100 | 100 | -------------- |
|  |  |  | sp. | ---------- | --------------- | ---- | ---- | -------------- |
|  | Annulatascales | |  |  |  |  |  |  |
|  |  | *Conlarium* | *duplumascospora* | BC1222 | JN936997.1 | 98 | 85 | -------------- |
|  |  | *Rhodoveronaea* | *varioseptata* | BC0645 | KF823603.1 | 99 | 82 | -------------- |
|  | Capnodiales | |  |  |  |  |  |  |
|  |  | *Acrocalymma* | *aquatica* | BC0359 | JX276951.1 | 83 | 83 | -------------- |
|  |  | *Cladosporium* | *cladosporioides* | BC1475 | MH714552.1 | 100 | 100 | -------------- |
|  |  |  | sp. | ---------- | --------------- | ---- | ---- | -------------- |
|  |  | *Passalora* | *brachycarpa* | BC1621 | GU214664.1 | 100 | 98 | -------------- |
|  |  | *Ramularia* | *coryli* | BC1480 | KX287391.1 | 100 | 98 | -------------- |
|  |  |  | *endophylla* | BC1635 | KP894243.1 | 98 | 98 | -------------- |
|  |  |  | *interstitiales* | BC1540 | KX287458.1 | 98 | 98 | -------------- |
|  |  |  | *rumicicola* | BC1601 | KX287503.1 | 100 | 96 | -------------- |
|  |  |  | sp. | ---------- | --------------- | ---- | ---- | -------------- |
|  |  | *Septoria* | *hyperici* | BC1571 | NR_147271.1 | 99 | 97 | -------------- |
|  |  | *Sphaerulina* | *berberidis* | BC1578 | LC206672.1 | 100 | 99 | -------------- |
|  |  | *Trichomerium* | *foliicola* | BC0315 | NR_144963.1 | 100 | 95 | -------------- |
|  | Chaetosphaeriales | |  |  |  |  |  |  |
|  |  | *Chaetosphaeria* | *chloroconia* | BC1533 | AF178542.1 | 98 | 99 | -------------- |
|  |  |  | *myriocarpa* | BC0320 | MH107883.1 | 98 | 99 | -------------- |
|  |  | *Chloridium* | *virescens* | BC1507 | EF029220.1 | 99 | 100 | -------------- |
|  |  |  | sp. | ---------- | --------------- | ---- | ---- | -------------- |
|  |  | *Codinaea* | *acaciae* | BC1318 | KY965397.1 | 100 | 96 | -------------- |
|  | Chaetothyriales | |  |  |  |  |  |  |
|  |  | *Capronia* | *dactylotricha* | BC1244 | NR_137136.1 | 91 | 86 | -------------- |
|  |  |  | *leucadendri* | BC0548 | NR_156212.1 | 98 | 98 | -------------- |
|  |  |  | *pilosella* | BC1634 | DQ826737.1 | 98 | 96 | -------------- |
|  |  |  | sp. | ---------- | --------------- | ---- | ---- | -------------- |
|  |  | *Cladophialophora* | *chaetospira* | BC1539 | KF359558.1 | 100 | 92 | -------------- |
|  |  |  | *potulentorum* | BC1241 | EU035410.1 | 99 | 90 | -------------- |
|  |  |  | sp. | ---------- | --------------- | ---- | ---- | -------------- |
|  |  | *Cyphellophora* | *gamsii* | BC1562 | NR_156306.1 | 100 | 95 | -------------- |
|  |  |  | *olivacea* | BC1633 | KX302010.1 | 95 | 98 | -------------- |
|  |  |  | *oxyspora* | BC0325 | MF196874.1 | 99 | 99 | -------------- |
|  |  |  | sp. | ---------- | --------------- | ---- | ---- | -------------- |
|  |  | *Exobasidium* | *otanianum* | BC1069 | AB180343.1 | 96 | 96 | -------------- |
|  |  | *Exophiala* | *moniliae* | BC1410 | HE605213.1 | 100 | 97 | -------------- |
|  |  |  | *xenobiotica* | BC1574 | KY434151.1 | 97 | 92 | -------------- |
|  |  |  | sp. | ---------- | --------------- | ---- | ---- | -------------- |
|  |  | *Fonsecaea* | *pedrosoi* | BC1176 | AB114131.1 | 98 | 98 | MH971241 |
|  |  |  | sp. (new) | BC1045 | JN999999.1 | 93 | 94 | MH971242 |
|  |  |  | sp. | ---------- | --------------- | ---- | ---- | -------------- |
|  |  | *Phialophora* | *americana* | BC1388 | U31840.1 | 99 | 99 | MH971243 |
|  |  |  | *sessilis* | BC1631 | GU981736.1 | 100 | 99 | MH971244 |
|  |  |  | sp. | ---------- | --------------- | ---- | ---- | -------------- |
|  |  | *Rhinocladiella* | *anceps* | BC0327 | AF050284.1 | 100 | 99 | MH971245 |
|  |  |  | *atrovirens* | BC0550 | AB091215.1 | 98 | 98 | MH971246 |
|  |  |  | *quercus* | BC1534 | NR_155728.1 | 97 | 99 | MH971247 |
|  | Coniochaetales | |  |  |  |  |  |  |
|  |  | *Coniochaeta* | *cephalothecoides* | BC0639 | KY064029.1 | 97 | 99 | -------------- |
|  |  |  | sp. | ---------- | --------------- | ---- | ---- | -------------- |
|  | Cordanales | |  |  |  |  |  |  |
|  |  | *Cordana* | *pauciseptata* | BC0927 | HE672147.1 | 93 | 99 | -------------- |
|  | Diaporthales | |  |  |  |  |  |  |
|  |  | *Cryptodiaporthe* | *hystrix* | BC0433 | KX776446.1 | 100 | 98 | -------------- |
|  |  | *Diaporthe* | *phaseolorum* | BC0083 | FJ441609.1 | 99 | 98 | -------------- |
|  |  | *Gnomonopsis* | sp. | ---------- | --------------- | ---- | ---- | -------------- |
|  |  | *Phaeoacremonium* | *iranianum* | BC1423 | KF764529.1 | 100 | 98 | -------------- |
|  |  |  | *mortoniae* | BC1555 | EU427312.1 | 82 | 99 | -------------- |
|  |  | *Togninia* | *minima* | BC0539 | KP083231.1 | 100 | 100 | -------------- |
|  | Dothideales | |  |  |  |  |  |  |
|  |  | *Aureobasidium* | *pullulans* | BC1444 | MF497401.1 | 100 | 100 | -------------- |
|  |  |  | sp. | ---------- | --------------- | ---- | ---- | -------------- |
|  |  | *Dothiora* | *sorbi* | BC0422 | KY929146.1 | 100 | 98 | -------------- |
|  |  |  | *pyrenophora* | BC1629 | KU728514.1 | 97 | 98 | -------------- |
|  | Eurotiales | |  |  |  |  |  |  |
|  |  | *Aspergillus* | *versicolor* | BC1564 | KU318416.1 | 98 | 99 | -------------- |
|  |  | *Paecilomyces* | *carneus* | BC1457 | HQ660442.1 | 100 | 99 | -------------- |
|  |  |  | *inflatus* | BC0451 | KU702692.1 | 99 | 99 | -------------- |
|  |  |  | *javanicus* | BC1300 | AB099944.1 | 100 | 100 | -------------- |
|  |  | *Penicillium* | *carneum* | BC1278 | NR_111551.1 | 100 | 100 | MH971248 |
|  |  |  | *chrysogenum* | BC1254 | MH778149.1 | 99 | 100 | MH971249 |
|  |  |  | *daleae* | BC1205 | MH854984.1 | 100 | 100 | MH971250 |
|  |  |  | *fellutanum* | BC1326 | HM469425.1 | 100 | 100 | MH971251 |
|  |  |  | *glabrum* | BC1095 | MF803957.1 | 100 | 100 | MH971252 |
|  |  |  | *herquei* | BC1214 | MF663569.1 | 100 | 100 | MH971253 |
|  |  |  | *nodositatum* | BC1031 | NR_103703.1 | 100 | 100 | MH971254 |
|  |  |  | *oxalicum* | BC1327 | MG733762.1 | 100 | 100 | MH971255 |
|  |  |  | *pancosmium* | BC1499 | MF803943.1 | 100 | 100 | MH971256 |
|  |  |  | *pinophilum* | BC0544 | EF488397.1 | 100 | 99 | MH971257 |
|  |  |  | *steckii* | BC0778 | MG554368.1 | 100 | 100 | MH971258 |
|  |  |  | *sumatraense* | BC1530 | JX140874.1 | 100 | 100 | MH971259 |
|  |  |  | sp. | ---------- | --------------- | ---- | ---- | -------------- |
|  |  | *Thysanophora* | *penicilloides* | BC1421 | JQ272462.1 | 99 | 100 | -------------- |
|  | Glomerellales | |  |  |  |  |  |  |
|  |  | *Glomerella* | *acutata* | BC1479 | JN697577.1 | 100 | 100 | -------------- |
|  | Helotiales | |  |  |  |  |  |  |
|  |  | *Cadophora* | *malorum* | BC0073 | DQ404350.1 | 100 | 98 | -------------- |
|  |  | *Catenulifera* | *brachyconia* | BC1570 | AB190384.1 | 100 | 97 | -------------- |
|  |  | *Hyalodendriella* | sp. | ---------- | --------------- | ---- | ---- | -------------- |
|  |  | *Hymenoscyphus* | *dehlii* | BC0543 | LC206621.1 | 94 | 94 | -------------- |
|  |  | *Hyphodiscus* | *hymeniophilus* | BC0581 | DQ227258.1 | 99 | 97 | -------------- |
|  |  | *Idriella* | *rara* | BC1306 | KC775737.1 | 91 | 98 | -------------- |
|  |  | *Leptodontidium* | *elatius* | BC1257 | AM981224.1 | 100 | 95 | -------------- |
|  |  |  | sp. | ---------- | --------------- | ---- | ---- | -------------- |
|  |  | *Leptodontium* | sp. | ---------- | --------------- | ---- | ---- | -------------- |
|  |  | *Pezicula* | *cinnamomea* | BC1553 | KR859145.1 | 98 | 83 | -------------- |
|  |  | *Pilidium* | *concavum* | BC1478 | MF776047.1 | 100 | 100 | -------------- |
|  |  | *Rhexocercosporidium* | sp. | ---------- | --------------- | ---- | ---- | -------------- |
|  |  | *Scytalidium* | *lignicola* | BC1391 | GQ272634.1 | 98 | 99 | -------------- |
|  |  |  | sp. | ---------- | --------------- | ---- | ---- | -------------- |
|  |  | *Synchaetomella* | *acerina* | BC1299 | NR_111811.1 | 99 | 93 | -------------- |
|  |  | *Xenopolyscytalum* | sp. | ---------- | --------------- | ---- | ---- | -------------- |
|  | Hypocreales | |  |  |  |  |  |  |
|  |  | *Acremonium* | *variecolor* | BC0031 | HE608648.1 | 99 | 99 | -------------- |
|  |  |  | sp. | ---------- | --------------- | ---- | ---- | -------------- |
|  |  | *Atractium* | *stilbaster* | BC1404 | KM231792.1 | 100 | 87 | -------------- |
|  |  | *Beauveria* | *brongniartii* | BC0062 | JX110373.1 | 99 | 99 | -------------- |
|  |  |  | *caledonica* | BC1250 | DQ350137.1 | 100 | 99 | -------------- |
|  |  | *Calcarisporium* | *arbuscula* | BC1182 | LC145810.1 | 100 | 99 | -------------- |
|  |  | *Clonostachys* | *rosea* | BC1297 | KX421414.1 | 99 | 99 | -------------- |
|  |  |  | sp. | ---------- | --------------- | ---- | ---- | -------------- |
|  |  | *Cordyceps* | *militaris* | BC1419 | KY407763.1 | 100 | 99 | -------------- |
|  |  |  | sp. | ---------- | --------------- | ---- | ---- | -------------- |
|  |  | *Cosmospora* | *butyri* | BC1580 | JQ070093.1 | 99 | 100 | MH971260 |
|  |  |  | *flavoviridis* | BC0394 | HQ897791.1 | 100 | 99 | MH971261 |
|  |  |  | *berkeleyana* | BC0428 | MH859583.1 | 99 | 96 | MH971262 |
|  |  |  | sp. | ---------- | --------------- | ---- | ---- | -------------- |
|  |  | *Cylindrium* | *elongatum* | BC1446 | KM231852.1 | 100 | 99 | -------------- |
|  |  | *Dialonectria* | sp. | ---------- | --------------- | ---- | ---- | -------------- |
|  |  | *Fusarium* | *equiseti* | BC1447 | GQ365157.1 | 100 | 100 | -------------- |
|  |  |  | *tricinctum* | BC1510 | JX045791.1 | 99 | 100 | -------------- |
|  |  |  | sp. | ---------- | --------------- | ---- | ---- | -------------- |
|  |  | *Fusicolla* | *melogramma* | BC0649 | NR_155096.1 | 99 | 99 | -------------- |
|  |  | *Geosmithia* | sp. | ---------- | --------------- | ---- | ---- | -------------- |
|  |  | *Gibberella* | sp. | ---------- | --------------- | ---- | ---- | -------------- |
|  |  | *Gliomastix* | *murorum* | BC0156 | AB540558.1 | 100 | 98 | -------------- |
|  |  | *Hirsutella* | sp. | ---------- | --------------- | ---- | ---- | -------------- |
|  |  | *Hypocrea* | *lutea* | BC1265 | AF359264.1 | 99 | 99 | -------------- |
|  |  |  | *pachybasioides* | BC0321 | AY240843.1 | 100 | 98 | -------------- |
|  |  | *Hypomyces* | *aurantius* | BC1020 | AB591044.1 | 100 | 98 | -------------- |
|  |  | *Isaria* | *farinosa* | BC0416 | MH191137.1 | 100 | 99 | -------------- |
|  |  |  | *fumosorosea* | BC0014 | FJ177460.1 | 100 | 99 | -------------- |
|  |  | *Lasionectria* | sp. | ---------- | --------------- | ---- | ---- | -------------- |
|  |  | *Lecanicillium* | *araneogenum* | BC0675 | KX845704.1 | 99 | 98 | -------------- |
|  |  |  | *attenuatum* | BC1503 | MH231313.1 | 100 | 100 | -------------- |
|  |  |  | *fungicola* | BC1022 | KX379184.1 | 99 | 100 | -------------- |
|  |  |  | *fusisporum* | BC1518 | AB378517.1 | 99 | 100 | -------------- |
|  |  |  | *psallotiae* | BC0015 | AB360367.1 | 99 | 99 | -------------- |
|  |  |  | *saksenae* | BC1455 | KY320616.1 | 100 | 97 | -------------- |
|  |  | *Mariannaea* | *elegans* | BC1489 | KX028787.1 | 99 | 99 | -------------- |
|  |  |  | *samuelsii* | BC1156 | KM231757.1 | 99 | 100 | -------------- |
|  |  | *Metacordyceps* | *chlamydosporia* | BC0618 | AB214654.1 | 100 | 99 | -------------- |
|  |  | *Metarhizium* | *anisopliae* | BC1203 | MG786739.1 | 100 | 100 | -------------- |
|  |  |  | *flavoviride* | BC1163 | KX380789.1 | 100 | 99 | -------------- |
|  |  | *Monocillium* | sp. | ---------- | --------------- | ---- | ---- | -------------- |
|  |  | *Nectria* | sp. | ---------- | --------------- | ---- | ---- | -------------- |
|  |  | *Niesslia* | *pulchriseta* | BC0033 | MG827040.1 | 96 | 99 | -------------- |
|  |  | *Pochonia* | *bulbillosa* | BC1436 | AB709835.1 | 100 | 99 | -------------- |
|  |  |  | *chlamydosporia* | BC1165 | AB713184.1 | 99 | 100 | -------------- |
|  |  |  | *suchlasporia* | BC0017 | AB214658.1 | 99 | 99 | -------------- |
|  |  | *Protocrea* | sp. | ---------- | --------------- | ---- | ---- | -------------- |
|  |  | *Pseudonectria* | sp. | ---------- | --------------- | ---- | ---- | -------------- |
|  |  | *Purpureocillium* | *lilacinum* | BC1070 | FJ765024.1 | 100 | 98 | -------------- |
|  |  | *Sepedonium* | *chalcipori* | BC1058 | KT946847.1 | 100 | 90 | -------------- |
|  |  | *Simplicillium* | *lamellicola* | BC1496 | KT004573.1 | 100 | 97 | -------------- |
|  |  |  | *lanosoniveum* | BC0027 | AB758126.1 | 99 | 99 | -------------- |
|  |  |  | sp. | ---------- | --------------- | ---- | ---- | -------------- |
|  |  | *Stilbella* | sp. | ---------- | --------------- | ---- | ---- | -------------- |
|  |  | *Tolypocladium* | *album* | BC1113 | MH137667.1 | 100 | 98 | -------------- |
|  |  |  | *inflatum* | BC0499 | KT693271.1 | 100 | 99 | -------------- |
|  |  | *Trichoderma* | *atroviride* | BC1215 | MF871528.1 | 100 | 100 | MH971263 |
|  |  |  | *harzianum* | BC1204 | MG132085.1 | 100 | 100 | MH971264 |
|  |  |  | *lixii* | BC1322 | MF782824.1 | 100 | 100 | MH971265 |
|  |  |  | *melanomagnum* | BC1271 | KU738454.1 | 100 | 100 | MH971266 |
|  |  |  | *pleuroticola* | BC1277 | MF687194.1 | 100 | 100 | MH971267 |
|  |  |  | *viride* | BC1433 | AJ230676.1 | 99 | 100 | MH971268 |
|  |  |  | sp. | ---------- | --------------- | ---- | ---- | -------------- |
|  |  | *Verticillium* | *fungicola* | BC0488 | FJ810136.1 | 100 | 99 | MH971269 |
|  |  |  | *insectorum* | BC1005 | AB214655.1 | 100 | 99 | MH971270 |
|  |  |  | *leptobactrum* | BC1341 | EF641871.1 | 99 | 99 | MH971271 |
|  |  |  | sp. | ---------- | --------------- | ---- | ---- | -------------- |
|  |  | *Xenoacremonium* | *falcatus* | BC1488 | MH062972.1 | 98 | 100 | -------------- |
|  | Mycosphaerellales | |  |  |  |  |  |  |
|  |  | *Mycosphaerella* | sp. | ---------- | --------------- | ---- | ---- | -------------- |
|  |  | *Ramichloridium* | sp. | ---------- | --------------- | ---- | ---- | -------------- |
|  | Onygenales | |  |  |  |  |  |  |
|  |  | *Oidiodendron* | sp. | ---------- | --------------- | ---- | ---- | -------------- |
|  | Ophiostomatales | |  |  |  |  |  |  |
|  |  | *Ophiostoma* | *dentifundum* | BC1140 | AY495435.1 | 97 | 93 | -------------- |
|  |  |  | *grandicarpum* | BC1638 | NR_147600.1 | 99 | 90 | -------------- |
|  |  | *Sporothrix* | *inflata* | BC0434 | AY495432.1 | 100 | 99 | -------------- |
|  |  |  | sp. | ---------- | --------------- | ---- | ---- | -------------- |
|  | Pleosporales | |  |  |  |  |  |  |
|  |  | *Coniothyrium* | *fuckelii* | BC0181 | AB665314.1 | 100 | 100 | -------------- |
|  |  | *Curvularia* | *geniculata* | BC1177 | MH517581.1 | 100 | 100 | -------------- |
|  |  |  | *senegalensis* | BC1151 | MH517581.1 | 100 | 100 | -------------- |
|  |  | *Dictyosporium* | *australiense* | BC0100 | DQ018092.1 | 97 | 98 | -------------- |
|  |  | *Helminthosporium* | *asterinum* | BC1248 | AF073917.1 | 99 | 99 | -------------- |
|  |  |  | *velutinum* | BC1385 | AB551948.1 | 98 | 99 | -------------- |
|  |  | *Leptosphaerulina* | *chartarum* | BC0086 | GU566269.1 | 100 | 99 | -------------- |
|  |  | *Lophiostoma* | sp. | ---------- | --------------- | ---- | ---- | -------------- |
|  |  | *Neocucurbitaria* | *vachellia* | BC1535 | NR_156363.1 | 100 | 95 | -------------- |
|  |  |  | *acerina* | BC1494 | MF795767.1 | 99 | 98 | -------------- |
|  |  | *Paraconiothyrium* | *brasiliense* | BC1485 | JF502455.1 | 99 | 99 | -------------- |
|  |  |  | *hawaiiense* | BC1456 | HM751092.1 | 100 | 99 | -------------- |
|  |  |  | sp. | ---------- | --------------- | ---- | ---- | -------------- |
|  |  | *Paraphaeosphaeria* | *neglecta* | BC1617 | JX496038.1 | 99 | 99 | -------------- |
|  |  | *Periconia* | *pseudobyssoides* | BC0452 | KC954161.1 | 100 | 99 | -------------- |
|  |  |  | sp. | ---------- | --------------- | ---- | ---- | -------------- |
|  |  | *Peyronellaea* | *glomerata* | BC0592 | MG832565.1 | 99 | 99 | -------------- |
|  |  | *Phaeosphaeria* | sp. | ---------- | --------------- | ---- | ---- | -------------- |
|  |  | *Phoma* | *bellidis* | BC1498 | JF502444.1 | 100 | 100 | -------------- |
|  |  |  | sp. | ---------- | --------------- | ---- | ---- | -------------- |
|  |  | *Preussia* | sp. | ---------- | --------------- | ---- | ---- | -------------- |
|  |  |  | sp. | ---------- | --------------- | ---- | ---- | -------------- |
|  |  | *Stagonosporopsis* | *cucurbitacearum* | BC1500 | LC168795.1 | 100 | 99 | -------------- |
|  |  | *Teichospora* | *quercus* | BC1412 | MH107920.1 | 98 | 96 | -------------- |
|  | Saccharomycetales | |  |  |  |  |  |  |
|  |  | *Candida* | *aglyptina* | BC1600 | FJ196775.1 | 76 | 98 | -------------- |
|  |  |  | *boleticola* | BC1585 | KY102001.1 | 100 | 99 | -------------- |
|  |  |  | sp. | ---------- | --------------- | ---- | ---- | -------------- |
|  |  | *Pichia* | *jadinii* | BC0555 | FJ865435.1 | 100 | 95 | -------------- |
|  |  |  | sp. | ---------- | --------------- | ---- | ---- | -------------- |
|  |  | *Yamadazyma* | *mexicana* | BC0590 | KY105944.1 | 100 | 99 | -------------- |
|  | Sordariales | |  |  |  |  |  |  |
|  |  | *Apodus* | *deciduus* | BC1170 | NR_145141.1 | 100 | 96 | -------------- |
|  |  | *Cercophora* | *sulphurella* | BC1619 | AY587913.1 | 99 | 96 | -------------- |
|  |  | *Chaetomium* | sp. | ---------- | --------------- | ---- | ---- | -------------- |
|  |  | *Fimetariella* | *rabenhorstii* | BC1129 | KX869958.1 | 98 | 98 | -------------- |
|  |  |  | sp. | ---------- | --------------- | ---- | ---- | -------------- |
|  |  | *Phialemonium* | *dimorphosporum* | BC0511 | FJ441614.1 | 100 | 98 | -------------- |
|  |  | *Spadicoides* | *bina* | BC0107 | JF340260.1 | 97 | 99 | -------------- |
|  | Trichosphaetiales | |  |  |  |  |  |  |
|  |  | *Nigrospora* | *oryzae* | BC1017 | HQ608152.1 | 100 | 100 | -------------- |
|  | Tubeufiales | |  |  |  |  |  |  |
|  |  | *Thaxteriellopsis* | *lignicola* | BC1594 | JN865207.1 | 99 | 78 | -------------- |
|  | Venturiales | |  |  |  |  |  |  |
|  |  | *Venturia* | *viennotii* | BC0500 | KF793787.1 | 97 | 96 | -------------- |
|  | Xylariales | |  |  |  |  |  |  |
|  |  | *Ascovirgaria* | *occulta* | BC0589 | AB740957.1 | 100 | 97 | -------------- |
|  |  | *Biscogniauxia* | *atropunctata* | BC0106 | JX507799.1 | 97 | 99 | -------------- |
|  |  |  | sp. | ---------- | --------------- | ---- | ---- | -------------- |
|  |  | *Diatrype* | *disciformis* | BC1565 | KR092795.1 | 99 | 99 | -------------- |
|  |  | *Eutypella* | *vitis* | BC1453 | KU320620.1 | 99 | 99 | -------------- |
|  |  | *Hansfordia* | *nebularis* | BC1173 | KF893290.1 | 99 | 100 | -------------- |
|  |  | *Hypoxylon* | *crocopeplum* | BC0178 | JN673047.1 | 99 | 99 | -------------- |
|  |  |  | *fendleri* | BC1400 | JN979417.1 | 100 | 92 | -------------- |
|  |  |  | *fuscum* | BC1159 | JN979424.1 | 100 | 96 | -------------- |
|  |  |  | *lenormandii* | BC1352 | KM610287.1 | 99 | 96 | -------------- |
|  |  |  | *perforatum* | BC1305 | JQ009308.1 | 100 | 99 | -------------- |
|  |  |  | *rubiginosum* | BC1438 | MH178694.1 | 91 | 99 | -------------- |
|  |  |  | *truncatum* | BC1079 | AF201716.2 | 98 | 99 | -------------- |
|  |  |  | sp. | ---------- | --------------- | ---- | ---- | -------------- |
|  |  | *Nemania* | *diffusa* | BC1521 | MH633932.1 | 99 | 91 | -------------- |
|  |  |  | *serpens* | BC1093 | KU683860.1 | 99 | 99 | -------------- |
|  |  |  | sp. | ---------- | --------------- | ---- | ---- | -------------- |
|  |  | *Spegazzinia* | sp. | ---------- | --------------- | ---- | ---- | -------------- |
|  |  | *Virgaria* | *nigra* | BC1406 | AB670716.1 | 100 | 98 | -------------- |
|  |  | *Xylaria* | *heliscus* | BC1355 | JQ761642.1 | 96 | 95 | MH971272 |
|  |  |  | sp. | ---------- | --------------- | ---- | ---- | -------------- |
|  | *incertae sedis* | |  |  |  |  |  |  |
|  |  | *Acrodontium* | *crateriforme* | BC1302 | KX287270.1 | 99 | 97 | -------------- |
|  |  | *Barbatosphaeria* | *dryina* | BC1060 | KM492892.1 | 92 | 96 | -------------- |
|  |  |  | *varioseptata* | BC1188 | NR_132089.1 | 92 | 97 | -------------- |
|  |  | *Geomyces* | sp. | ---------- | --------------- | ---- | ---- | -------------- |
|  |  | *Leohumicola* | sp. | ---------- | --------------- | ---- | ---- | -------------- |
|  |  | *Pseudogymnoascus* | sp. | ---------- | --------------- | ---- | ---- | -------------- |
|  |  | *Ramimonilia* | *apicalis* | BC1546 | NR_144959.1 | 89 | 93 | -------------- |
|  |  | *Veronaea* | *japonica* | BC1623 | NR_111277.1 | 98 | 97 | -------------- |
|  |  | *Xylomelasma* | sp. | ---------- | --------------- | ---- | ---- | -------------- |
| Basidiomycota | | |  |  |  |  |  |  |
|  | Agaricales | |  |  |  |  |  |  |
|  |  | *Schizophyllum* | *commune* | BC1449 | MF554593.1 | 100 | 99 | -------------- |
|  | Cantharellales | |  |  |  |  |  |  |
|  |  | *Sistotrema* | *brinkmanii* | BC1078 | KX527870.1 | 99 | 99 | -------------- |
|  |  |  | *oblongisporum* | BC1508 | KP814309.1 | 99 | 99 | -------------- |
|  |  | *Tulasnella* | sp. | ---------- | --------------- | ---- | ---- | -------------- |
|  | Dacrymycetales | |  |  |  |  |  |  |
|  |  | *Calocera* | sp. | ---------- | --------------- | ---- | ---- | -------------- |
|  |  | *Dacrymyces* | *australis* | BC1195 | MH858261.1 | 96 | 84 | -------------- |
|  | Exobasidiales | |  |  |  |  |  |  |
|  |  | *Meira* | sp. | ---------- | --------------- | ---- | ---- | -------------- |
|  | Hymenochaetales | |  |  |  |  |  |  |
|  |  | *Hydnochaete* | *olivacea* | BC1560 | KJ140712.1 | 100 | 82 | -------------- |
|  |  | *Peniophorella* | *pubera* | BC1138 | KP715565.1 | 97 | 84 | -------------- |
|  | Malasseziales | |  |  |  |  |  |  |
|  |  | *Malassezia* | sp. | ---------- | --------------- | ---- | ---- | -------------- |
|  | Microbotryales | |  |  |  |  |  |  |
|  |  | *Curvibasidium* | *cygneicollum* | BC1606 | KY102973.1 | 98 | 98 | -------------- |
|  |  |  | *pallidicorallinum* | BC0542 | JX188149.1 | 100 | 99 | -------------- |
|  | Microstromatales | |  |  |  |  |  |  |
|  |  | *Microstroma* | *juglandis* | BC1607 | EU069498.1 | 95 | 90 | -------------- |
|  | Polyporales | |  |  |  |  |  |  |
|  |  | *Bjerkandera* | *adusta* | BC1316 | MF161298.1 | 100 | 99 | -------------- |
|  |  | *Ceriporia* | *lacerata* | BC1158 | KP135024.1 | 99 | 100 | -------------- |
|  |  |  | sp. | ---------- | --------------- | ---- | ---- | -------------- |
|  |  | *Ceriporiopsis* | *gilvescens* | BC0174 | KY948761.1 | 100 | 100 | -------------- |
|  |  | *Gloeoporus* | *pannocinctus* | BC0042 | JQ673102.1 | 100 | 100 | -------------- |
|  |  | *Irpex* | *lacteus* | BC1038 | JX290579.1 | 100 | 99 | -------------- |
|  |  |  | sp. | ---------- | --------------- | ---- | ---- | -------------- |
|  |  | *Junghuhnia* | *nitida* | BC0528 | KP135323.1 | 97 | 99 | -------------- |
|  |  | *Phanerochaete* | *cumulodentata* | BC0709 | KP994373.1 | 100 | 100 | MH971273 |
|  |  |  | *sordida* | BC0691 | KP135074.1 | 100 | 100 | MH971274 |
|  |  |  | sp. | ---------- | --------------- | ---- | ---- | -------------- |
|  |  | *Phlebia* | *acerina* | BC1331 | KP135373.1 | 100 | 99 | -------------- |
|  |  |  | *fuscoatra* | BC1014 | KY948754.1 | 99 | 97 | -------------- |
|  |  |  | *lividina* | BC0629 | KY948756.1 | 100 | 99 | -------------- |
|  |  |  | *subserialis* | BC0054 | HQ607954.1 | 100 | 99 | -------------- |
|  |  |  | *tremellosa* | BC0184 | GU062266.1 | 99 | 99 | -------------- |
|  |  | *Phlebiopsis* | *flavidoalba* | BC1426 | KP135401.1 | 100 | 100 | -------------- |
|  |  |  | *gigantea* | BC1049 | KX028786.1 | 100 | 100 | -------------- |
|  |  | *Scopuloides* | *rimosa* | BC1046 | KP135348.1 | 99 | 100 | -------------- |
|  |  | *Trametopsis* | *cervina* | BC1143 | JX463662.1 | 99 | 99 | -------------- |
|  | Russulales | |  |  |  |  |  |  |
|  |  | *Peniophora* | *pithya* | BC1467 | MH857635.1 | 100 | 95 | -------------- |
|  |  |  | sp. | ---------- | --------------- | ---- | ---- | -------------- |
|  |  | *Stereum* | *complicatum* | BC1428 | MF161283.1 | 98 | 100 | -------------- |
|  |  |  | sp. | ---------- | --------------- | ---- | ---- | -------------- |
|  | Sporidiobolales | |  |  |  |  |  |  |
|  |  | *Rhodotorula* | *eucalyptica* | BC1579 | EU075186.1 | 99 | 87 | -------------- |
|  |  |  | sp. | ---------- | --------------- | ---- | ---- | -------------- |
|  | Tremellales | |  |  |  |  |  |  |
|  |  | *Cryptococcus* | *dimennae* | BC0392 | KM246197.1 | 98 | 98 | -------------- |
|  |  |  | sp. | ---------- | --------------- | ---- | ---- | -------------- |
|  |  | *Kockovaella* | *fuzhouensis* | BC1536 | KY103850.1 | 97 | 93 | -------------- |
|  |  |  | *sichuanensis* | BC1542 | KY103859.1 | 100 | 99 | -------------- |
|  |  | *Tremella* | *subalpina* | BC1424 | NR_155908.1 | 91 | 85 | -------------- |
|  | *incertae sedis* | |  |  |  |  |  |  |
|  |  | *Acaromyces* | *ingoldii* | BC1417 | HM595575.1 | 100 | 100 | -------------- |
|  |  | *Hamamotoa* | *singularis* | BC1050 | AF444600.1 | 100 | 87 | -------------- |
| Mucoromycota | | |  |  |  |  |  |  |
|  | Mortierellales | |  |  |  |  |  |  |
|  |  | *Mortierella* | aff. *ambigua* | BC1065 | JX976067.1 | 100 | 92 | MH971275 |
|  |  |  | sp. | ---------- | --------------- | ---- | ---- | -------------- |
|  | Mucorales | |  |  |  |  |  |  |
|  |  | aff. *Apophysomyces* | sp. | BC1021 | NR_137035.1 | 91 | 78 | MH971276 |
|  |  | *Backusella* | *circina* | BC1287 | JQ979443.1 | 100 | 94 | -------------- |
|  |  |  | *recurva* | BC1267 | JN206265.1 | 99 | 98 | -------------- |
|  |  | *Cunninghamella* | *elegans* | BC1044 | MH857146.1 | 99 | 99 | -------------- |
|  |  | *Mucor* | *abundans* | BC1003 | MH855716.1 | 99 | 99 | MH971277 |
|  |  |  | *circinelloides* | BC1285 | MH856522.1 | 99 | 100 | MH971278 |
|  |  |  | *fragilis* | BC1294 | KY047150.1 | 100 | 99 | MH971279 |
|  |  |  | *genevensis* | BC1002 | JN206046.1 | 97 | 98 | MH971280 |
|  |  |  | *luteus* | BC1107 | MH859851.1 | 98 | 98 | MH971281 |
|  |  |  | *mucedo* | BC1056 | JQ319046.1 | 100 | 98 | MH971282 |
|  |  |  | *racemosus* | BC1295 | MF356581.1 | 100 | 99 | MH971283 |
|  |  |  | sp. | ---------- | --------------- | ---- | ---- | -------------- |
|  | Umbelopsidales | |  |  |  |  |  |  |
|  |  | *Umbelopsis* | *angularis* | BC0529 | MH859191.1 | 99 | 99 | MH971284 |
|  |  |  | *isabellina* | BC1080 | KM044070.1 | 100 | 99 | MH971285 |
|  |  |  | *ramanniana* | BC1097 | MH864647.1 | 100 | 100 | MH971286 |
|  |  |  | sp. | ---------- | --------------- | ---- | ---- | -------------- |
| Zoopagomycota | | |  |  |  |  |  |  |
|  | Entomophthorales | |  |  |  |  |  |  |
|  |  | *Basidiobolus* | *ranarum* | BC1232 | EF392530.1 | 100 | 99 | -------------- |
